# *LINC00205* acts as a multivalent scaffold promoting FUS^P525L^ recruitment in Amyotrophic Lateral Sclerosis stress granules

**DOI:** 10.64898/2026.04.08.715854

**Authors:** Jessica Rea, Gaia Stortini, Tiziana Santini, Adriano Setti, Marta Bernardi, Pierpaolo Cantisani, Letizia Fucci, Davide Mariani, Paolo Tollis, Erika Vitiello, Irene Bozzoni

## Abstract

Stress granules (SG) are dynamic, membrane-less ribonucleoprotein assemblies that orchestrate cellular stress responses and rapidly disassemble upon stress relief. In Amyotrophic Lateral Sclerosis (ALS), mutations in RNA-binding proteins such as Fused in Sarcoma (FUS) impair SG dynamics, promoting the formation of aberrant and persistent granules. Although long non-coding RNAs (lncRNAs) are emerging as regulators of ribonucleoprotein organization, their mechanistic role in SG architecture and pathological remodeling remains largely unexplored. Here, we identify *LINC00205* as a critical RNA regulator of pathological SG dynamics in FUS^P525L^-associated ALS.

Using Neuroblastoma cells and human iPSC-derived Motor Neurons (MN), we show that *LINC00205* is enriched in SG upon oxidative stress and directly interacts with mutant FUS^P525L^. Knock-out of *LINC00205* selectively reduces the formation of FUS^P525L^-containing SG and restores physiological SG disassembly kinetics, without affecting normal SG or FUS expression levels. Mechanistically, *LINC00205* acts as a multivalent RNA scaffold, directly binding mRNAs specifically enriched in pathological SG, such as *PLCXD3* and *PIK3CA*, as well as the RNA helicase DHX36, which is preferentially associated with FUS^P525L^-containing SG. We demonstrate that *LINC00205* promotes their specific recruitment into pathological SG, thereby contributing to the aberrant features of these assemblies.

Together, our findings uncover an active lncRNA-driven mechanism that shapes the molecular composition of aberrant ALS-related SG and regulates their persistence, establishing lncRNAs as key organizers of RNA-protein assemblies under stress and providing a conceptual framework for modulating pathological condensates.

## INTRODUCTION

Cells spatially organize macromolecules such as proteins and RNAs in membrane-less organelles, creating transient functional condensates that mediate essential biological processes, such as cytoprotection [1–3]. The formation and stability of these organelles are mediated by liquid-liquid phase separation (LLPS) events [4–6]. LLPS is environmentally regulated and mediates key aspects of the cellular stress response: the formation of reversible, phase-separated structures allows cells to temporarily sequester proteins and RNAs and to resume function rapidly once stress subsides [3,7–9].

Among the cytoplasmic membrane-less organelles, stress granules (SG) have critical roles in mRNA metabolism and translational control [10,11]. Cells assemble SG rapidly in response to a variety of stress conditions, leading to the suppression of global protein synthesis while promoting translation of stress-responsive transcripts. When stress ends, SG disperse and stored mRNAs re-enter translation [9,12–14].

Proteomic analyses indicate that about 50% of SG components are RNA-binding proteins (RBPs) [15]. These RBPs help drive phase separation through intrinsically disordered, low-complexity domains (LCDs) and engage multivalent interactions with other macromolecules [16,17].

The dynamism of SG is lost in neurodegenerative diseases, including Amyotrophic Lateral Sclerosis (ALS). Under sustained stress, RNA-binding proteins accumulate in SG, leading to their maturation into insoluble aggregates [18–20]. A typical example is represented by Fused in Sarcoma (FUS), an ALS-associated RBP that mislocalizes to the cytoplasm where it can undergo phase separation leading to the formation of pathological aggregates [21–23]. Mutations in FUS account for about 4% of familiar ALS cases and cluster within the C-terminal region [24,25]. In particular, the proline-to-lysine substitution at position 525 (P525L) is frequently linked to juvenile ALS and is reported in roughly three out of ten juvenile cases, typically with onset before twenty-five years of age [26].

Some long non-coding RNAs (lncRNAs) show stress-dependent regulation and relocalization, and increasing local RNA concentration can nucleate ribonucleoprotein (RNP) condensates under stress [27,28]. In neurons, lncRNAs add a critical regulatory layer in the biology of biomolecular condensates by acting as multivalent scaffolds that bind multiple RBPs and RNAs, sustaining a dynamic interaction network [29,30]. *NEAT1* exemplifies this principle in paraspeckles and contributes to stress responses, while mislocalized TAR DNA-binding protein 43 (TDP-43) and FUS can disrupt *NEAT1* processing and compromise organelle function [31–33]. SG display selective RNA content that is remodeled in ALS, with mutant FUS contributing to this shift [23], which implies an active role of RNA species, including lncRNAs, in characterizing condensate composition and properties [34–36].

LncRNAs are known to act as architects during the formation of membrane-less organelles. In this context, our aim is to explore their active role in the pathological remodeling of SG in ALS, a question that currently remains unexplored. In this work, we identify *LINC00205* as a regulator of pathological SG in FUS^P525L^-associated ALS. We demonstrate that it recruits FUS^P525L^ and other specific RNA and protein components into aberrant SG, controlling their disassembly kinetics without affecting physiological SG or FUS levels. These results shed light on a novel lncRNA-mediated mechanism controlling aberrant SG in FUS^P525L^-associated ALS and establish lncRNAs as attractive targets to modulate pathological condensates in neurodegenerative diseases.

## MATERIALS AND METHODS

### SK-N-BE cell culture, maintenance and treatment

SK-N-BE (ATCC) cells engineered to express Doxycycline (DOX)-inducible FLAG-FUS construct, either wild-type (FUS^WT^) or carrying the P525L mutation (FUS^P525L^), as described in [37], were used. To induce construct expression, cells were exposed to 50 ng/ml DOX (Sigma-Aldrich, #D9891) for 24h. Cells were maintained in adherent culture in RPMI-1640 medium (Sigma-Aldrich, cat#R6504) supplemented with 10% fetal bovine serum (Sigma-Aldrich, cat#F2442), 1:100 GlutaMAX™ (ThermoFisher cat#35050061), 1:100 Sodium Pyruvate (Sigma-Aldrich, cat#P5280), and 1:100 Penicillin/Streptomycin (Sigma-Aldrich cat#P0781). For subculturing, cells were washed with Dulbecco’s Phosphate Buffered Saline w/o MgCl_2_ (PBS w/o) (Sigma-Aldrich cat#D8537) and dissociated using Trypsin solution from porcine pancreas (Sigma-Aldrich cat#T4549). Cells were frozen in culture medium supplemented with 20% FBS and 10% Dimethyl Sulfoxide (DMSO, Sigma Aldrich) and quickly thawed when needed in a 37°C water bath. Acute oxidative stress was induced by treating cells with 0.5mM Sodium Arsenite (ARS, Sigma-Aldrich, #106277) for 1 h.

### FUS^WT/P525L^ hiPSC culture and differentiation into spinal Motor Neurons

Human induced pluripotent stem cells (hiPSC) were used in this study. Specifically, two hiPSC lines expressing both FUS^WT^ and FUS^P525L^ were adopted, derived by [38]. hiPSC were maintained in 0,1 mg/ml Geltrex^TM^-coated (Gibco cat#A1413201) plates in NutriStem® hiPSC XF medium (Sartorius, #05-100-1 A) supplemented with 1:1000 Penicillin/Streptomycin (Sigma-Aldrich cat#P0781). For subculturing, cells were treated with 1X Dispase II (Gibco cat#17105041) for colony dissociation, or Accutase® (Thermo Fisher Scientific, #00-4555-56), for single cell dissociation. hiPSC were induced to differentiate into spinal Motor Neurons (MN) following the methods described in [38]. Acute oxidative stress was induced by treating cells with 0.5mM Sodium Arsenite (ARS) (Sigma-Aldrich, #106277) for 1 h.

### Immunofluorescence

Cells were cultured on glass coverslips pre-coated with 0,1 mg/ml Geltrex^TM^-coated (Gibco cat#A141320. Fixation was carried out in 4% paraformaldehyde (Electron Microscopy Sciences, cat#157-8) diluted in Dulbecco’s Phosphate Buffered Saline with MgCl_2_ and CaCl_2_ (complete PBS) (Sigma- Aldrich, cat# D86622), for 20 min at 4°C. After washing with complete PBS, cells were permeabilized using 0.2% Triton X-100 (PBS-T) (Sigma-Aldrich, cat#9036-19-59) diluted in complete PBS and incubated for 15 min at room temperature (RT). Blocking was carried out with 3% Goat Serum in complete PBS for 20 min at RT in a humidified chamber. Primary antibodies were incubated overnight at 4°C diluted in blocking solution. The following antibodies were used: 1:150 anti-HuR (Santa Cruz biotechnology, sc-5261), 1:250 anti-DHX36 (Proteintech, 13159-1-AP), 1:300 anti-FUS (Santa Cruz biotechnology, sc-47711), 1:300 anti-G3BP1(Abcam, car# EPR13986(B)), 1:300 anti-TUB (Sigma-Aldrich, cat#AB9354); 1:200 anti-TIAR (BD Transduction, cat#610352). After two washes with PBS-T for 3 minutes, secondary antibodies were incubated for 1 h at room temperature in blocking solution: anti-mouse Alexa Fluor® 647 (Thermo Fisher, cat#A-31571), anti-rabbit Alexa Fluor® 488 (Thermo Fisher, cat#A-11034), anti-chicken Alexa Fluor® 594 (Thermo Fisher, cat#A-11001); all 1:300. Finally, the cells were stained with DAPI (3 µg/mL in PBS-T) for 5 min. After a final wash with PBS-T, coverslips were mounted using ProLong™ Diamond Antifade Mountant (Invitrogen, cat#P36961). Slides were cured in the dark at room temperature for 48h and stored at −20°C until imaging.

### Immunofluorescence coupled with single molecule fluorescence in situ hybridization

To visualize *LINC00205*, *GAPDH*, *PLCXD3*, and *PIK3CA* mRNAs, a multiplexed hybridization chain reaction fluorescence in situ hybridization protocol (HCR-FISH) was performed in combination with immunofluorescence, as described [39,40]. The synthetic DNA oligonucleotides used for HCR-FISH were designed as described in [40] and are listed in Supplementary Table 1.

### Confocal microscopes

Samples were imaged with a 60X oil objective using: i) inverted Olympus IX73 microscope/X-LIGHT V3 spinning disk integrated and a Prime BSI Express Scientific CMOS camera; ii) Olympus iX83 FluoView1200 laser scanning confocal microscope; iii) Nikon ECLIPSE Ti2-E microscope/X-Light V3 spinning disk integrated equipped with Teledyne Photometrics sCMOS Kinetix camera.

### SG count/recovery analyses

Images were analyzed using the open-source software FIJI/ImageJ. Briefly, after background subtraction, G3BP1 and cytoplasmic FUS spots were quantified using the 3D Object Counter. To quantify cytoplasmic FUS signal, the FUS staining localized in the nucleus was subtracted by superimposing a DAPI mask before the quantification. To evaluate recovery after stress removal, cells containing at least one G3BP1-positive SG were counted and related to the total cells in each microscope field.

### Colocalization analyses

To measure RNA enrichment in SG, quantification was performed on Maximum Intensity Projection (MIP) of confocal Z-stack. After background removal, smFISH spots were detected using Find Maxima. SmFISH signals were quantified before and after G3BP1 mask subtraction to calculate the percentage of spots colocalizing with G3BP1.

To evaluate DHX36 and HuR recruitment within SG, TIAR was used as SG marker for colocalization with DHX36, while G3BP1 was used as SG marker for colocalization with HuR. After background removal, Pearson’s correlation coefficients were calculated on MIP using the FIJI/ImageJ JaCoP plugin.

### *LINC00205* knock-out through CRISPR/Cas9 engineering

Two single-guide RNAs (sgRNAs) targeting *LINC00205* were designed using Benchling (https://www.benchling.com/) design tool (GUIDE1: 5’-GGGGCTCCTGAGGGTCCGCG-3’; GUIDE2: 5’-TGAGGGTCCGCGAGGCCGGG-3’) and cloned into the pX333 plasmid encoding wild-type Cas9 (PX333-sgRNAs) [41]. The *LINC00205* transcription start site was identified via Zenbu (FANTOM5) to define Cas9 target regions. HR110PA (System Biosciences) was used as a backbone to create the donor vector (DONOR). A Poly-adenylation sequence (PAS) was cloned into the DONOR followed by a Neomycin resistance cassette using In-Fusion® HD Cloning Plus Kit (Cat. #638910). Two homology arms (HA) HA1 and HA2, with a length of 800 nt were amplified by PCR from iPSC gDNA (Kapa HiFi, Takara Bio). HA1 was cloned upstream of the PAS, and HA2 was cloned downstream of the PAS. FUS^WT^ and FUS^P525L^ hiPSC were transfected on Geltrex^TM^-coated dishes using the Neon Transfection System (Life Technologies; 100 µL tips, 1200 V, 30 ms, 1 pulse) in R buffer. Selection was performed with 800 µg/mL G418 for 5 days. Control lines were maintained in parallel without transfection. Single *LINC00205* knock-out (KO) clones were isolated and genotyped. Oligonucleotides used to assess *LINC00205* KO are listed in Supplementary Table 1.

### RNA extraction and analysis

RNA from immunoprecipitation (IP) and pull-down (PD) experiments was extracted using the QIAGEN RNeasy® Mini Kit (QIAGEN, #74104), whereas RNA from total cellular extracts was isolated with the Direct-zol RNA Miniprep Kit (Zymo Research, #R2053), following the manufacturers’ protocol. RNA obtained from IP and PD experiments was reverse-transcribed into cDNA using VILO cDNA Synthesis Kit (Invitrogen, cat#11754050), while RNA from total cellular extracts was reverse transcribed using the Takara PrimeScript™ RT Reagent Kit (Takara-bio, cat#RR036b). For samples requiring removal of residual DNA (e.g., native PD RNAs with biotinylated DNA probes), an extra DNase I (Thermo Fisher, #EN0521) treatment was performed prior to reverse transcription. Quantitative real-time PCR (qRT-PCR) was carried out using PowerUp™ SYBR™ Green Master Mix (Thermo Fisher, #A25742) on a QuantStudio 3 Real-Time PCR system (Thermo Fisher Scientific). Oligonucleotides used for qRT-PCR assays are listed in Supplementary Table 1.

### Protein extraction and western blot

Whole-cell protein extracts were prepared from hiPSC-derived MN and SK-N-BE cells lysed in RIPA buffer (50 mM Tris–HCl (pH 8), 150 mM EDTA (Sigma Life Science, cat#E5134-1KG), 150 mM NaCl (Sigma-Aldrich, cat#S9888-5KG), 50 mM NaF (Sigma-Aldrich 7681-49-4), 10% glycerol (Sigma-Aldrich, cat#G2289), 1.5 mM MgCl2 (Electron Microscopy Sciences, cat#18010), 1% Triton X-100 (Sigma-Aldrich, cat#9036-19-59). Proteins were separated using the NuPAGE™ System (Invitrogen, #EI0002) on 4–12% Bis-Tris gels (Invitrogen, #NP0321BOX) and transferred to Amersham Protran 0.45 µm nitrocellulose membranes (GE Healthcare, #10600002) using the Mini Trans-Blot® Cell (Bio-Rad, #1703810). Membranes were incubated with the following antibodies: anti-GAPDH (Abcam, ab8245) 1:1000, anti- α Actinin (Santa Cruz, sc-390205) 1:5000, anti-Flag M2-Peroxidase (HRP) (Sigma-Aldrich, #A8592) 1:2500, anti-HuR (Santa Cruz biotechnology, sc-5261) 1:1000, anti-DHX36 (Proteintech, 13159-1-AP) 1:1000, anti-FUS (Santa Cruz biotechnology, sc-47711) 1:1000.

Protein signals were detected using: WesternBright ECL (Advansta, #K-12045-D50) acquired with a ChemiDoc XRS+ Molecular Imager (Bio-Rad) or with Fluorescent secondary antibodies StarBright Blue 520 Goat Anti-Rabbit IgG (Sigma-Aldrich, cat# 12005869) and StarBright Blue 700 Goat Anti-Mouse IgG (Sigma-Aldrich, cat#12004158), acquired with a ChemiDoc MP Molecular Imager (Bio-Rad). Quantification was performed using Image Lab Software (v6.1.0).

### CLIP assays

SK-N-BE were crosslinked using 460 nm UV light at 4000 × 100 μJ/cm² in complete PBS and lysed in 500 µl Complete Lysis Buffer (20 mM Tris-HCl (Sigma-Aldrich cat#11814273001, cat#30721-1L) pH 7.5, 100 mM NaCl (Sigma-Aldrich, cat#S9888-5KG), 0.5 mM EDTA (Sigma Life Science, cat#E5134-1KG), 0,5% 0,5% NP-40 (Sigma-Aldrich, cat#85124), 0.1% SDS (AppliChem, cat#A0675,0500) supplemented with 1X PIC (Roche, cat#11873580001), 1mM Dithiothreitol (DTT) (Roche, cat#71717923), 0.16 U/µL RiboLock RNase inhibitor (ThermoFisher, cat# EO038SKB011)). Lysates were clarified by centrifugation at 16,000 × g for 10 min at 4 °C and quantified via Bradford assay. For DHX36 and HuR CLIP assays, 50 μL of Dynabeads^TM^ Protein A (Thermo Fisher Scientific, cat#10002D) or Protein G (Thermo Fisher Scientific, cat#10004D), respectively, were washed twice with PBS w/o (Sigma-Aldrich cat#D8537) supplemented with 0.02% Tween® 20 (Sigma-Aldrich, cat# 1379) (PBS-T). Beads were conjugated with 5 µg of antibody (IP or IgG) in PBS-T by incubation on rotating wheel for 2.5 h at room temperature. The following antibodies were used for IPs: anti-DHX36 (Proteintech, 13159-1-AP), anti-HuR (Santa Cruz biotechnology, sc-5261). For FLAG-FUS^P525L^ CLIP, anti-FLAG beads (Sigma, # M8823) were used. Beads were washed twice with PBS-T, mixed with 500–1000 µg extract, adjusted to 1ml with complete lysis buffer and incubated overnight at 4 °C on rotating wheel. Input (INP) samples were prepared with 10% of the total extract and adjusted to 100 µl with complete lysis buffer. Beads were washed once with Wash Buffer (50mM Tris-HCl (Sigma-Aldrich cat#11814273001, cat#30721-1L), 150 mM NaCl (Sigma-Aldrich, cat#S9888-5KG), 1mM MgCl2 (Electron Microscopy Sciences, cat#18010), 0,50% NP-40 (Sigma-Aldrich, cat#85124)) for 2 minutes, and three times with High Salt Wash Buffer (50mM Tris-HCl (Sigma-Aldrich cat#11814273001, cat#30721-1L), 500 mM NaCl (Sigma-Aldrich, cat#S9888-5KG), 1mM MgCl2 (Electron Microscopy Sciences, cat#18010), 0,50% NP-40 (Sigma-Aldrich, cat#85124)) for 30 seconds. All wash incubations were done on a rotating wheel. Beads were resuspended in 100 µl complete lysis buffer and split ¾ for RNA extraction and ¼ for protein analysis. RNA fractions were treated with a mix composed by 99,5 µL of Proteinase K Buffer (200 mM Tris-HCl (Sigma-Aldrich cat#11814273001, cat#30721-1L), 200 mM NaCl (Sigma-Aldrich, cat#S9888-5KG), 25 mM EDTA (Sigma Life Science, cat#E5134-1KG), 2% SDS 20% (AppliChem, cat#A0675,0500)), 0,5 µL RiboLock RNase inhibitor (ThermoFisher, cat# EO038SKB011), 10 µL Proteinase K, recombinant PCR Grade (Roche, cat#03115828001), 15 µL complete lysis buffer. Samples were incubated for 1 h at 70 °C and 800 RPM, and 750 µL of Trizol lysis reagent were added. Protein fractions were mixed with 10 µL 4× Laemmli Sample Buffer (Bio-Rad, cat#161-0747) and 4 µL 1 mM DTT (Roche, cat#71717923). Samples were heated at 90 °C for 5 minutes at 1000 RPM, and beads were then removed.

### Native RNA Pull-Down

5 × 10^7^ SK-N-BE cells were resuspended in 1ml of Lysis buffer (50mM Tris-HCl (Sigma-Aldrich cat#11814273001, cat#30721-1L), 150 mM NaCl (Sigma-Aldrich, cat#S9888-5KG), 3mM 1mM MgCl2 (Electron Microscopy Sciences, cat#18010), 2mM EDTA (Sigma Life Science, cat#E5134-1KG), 0,5% NP-40 (Sigma-Aldrich, cat#85124)) complemented with 1X PIC (Roche, cat#11873580001), 1mM DTT (Roche, cat#71717923), 0.16 U/µL RiboLock RNase inhibitor (ThermoFisher, cat# EO038SKB011). Lysates were centrifuged at 15.000 × g for 15 minutes at 4°C. The supernatant was collected for quantification via Bradford assay. 1 mg of extract was used for each pull-down (PD) condition (*LINC00205* or *LACZ)* and then adjusted to 1ml with complete lysis buffer. To each sample, 600 µL hybridization buffer (100mM Tris-HCl (Sigma-Aldrich cat#11814273001, cat#30721-1L) (Sigma-Aldrich cat#11814273001, cat#30721-1L) 300 mM NaCl (Sigma-Aldrich, cat#S9888-5KG), 1mM MgCl_2_ (Electron Microscopy Sciences, cat#18010), 10mM EDTA (Sigma Life Science, cat#E5134-1KG), 15% Formamide (Sigma-Aldrich, cat# 47671), 0.2% SDS (AppliChem, cat#A0675,0500), 0,5% NP-40 (Sigma-Aldrich, cat#85124)) complemented with 1X PIC (Roche, cat#11873580001), 1mM DTT (Roche, cat#71717923), 0.16 U/µL RiboLock RNase inhibitor (ThermoFisher, cat# EO038SKB011), 105.55 µL 2,5% Dextran sulfate sodium salt (Sigma-Aldrich, cat# D8906), and 15-30 µL 10 pmol/µL 5′-biotinylated 20-nt antisense DNA probes (listed in Supplementary Table 1), preheated at 80 °C for 3 min, were added. For the *LINC00205* PD assay followed by RNA-seq, ODD and EVEN probe sets were used, whereas a pooled probe mix was employed for *LINC00205* PD validation. In parallel, an INP sample was prepared with 10% of the total extract in 100 µL complete lysis buffer and supplemented with 60 µL complete hybridization buffer. The extracts were incubated for 4 hours on a rotating wheel at 4°C. Streptavidin MagnaSphere® paramagnetic particles (Promega, cat#Z5481) (150–300 µL) were washed twice with hybridization buffer, resuspended in 150–300 µL hybridization buffer, added to samples, and incubated 1h at room temperature on a rotating wheel. Beads were washed 4 times with hybridization buffer (3 min per wash at 4 °C on a rotating wheel), and then 500 µL Trizol lysis reagent was added to beads and INP for downstream RNA extraction.

### Preparation of RNA libraries and RNA-sequencing

RNA libraries were prepared using the Illumina Stranded Total RNA Prep with Ribo-Zero^TM^ Plus kit and sequenced on an Illumina NovaSeq 6000 platform in paired-end mode, generating reads of 100 bases. Read quality was assessed with FastQC v0.11.9, which revealed a transient reduction in quality at the first base, residual contamination consistent with incomplete ribosomal RNA removal by Ribo-Zero Plus, and the presence of Illumina adapter sequences. Adapter trimming and removal of low-quality nucleotides were performed using cutadapt v3.2 [42] with: *-u 1 -U 1 --trim-n --nextseq-trim=20 -m 18* and Trimmomatic software v0.39 [43] with PE mode and the following parameters; *ILLUMINACLIP:adapter_path:2:30:10:8:true LEADING:3 TRAILING:3 SLIDINGWINDOW:4:20 MINLEN:18*. To eliminate remaining ribosomal RNA, trimmed reads were aligned to an rRNA-only reference (Supplementary Table 2) using Bowtie2 v2.4.2 [44], and only unmapped reads were retained. These filtered reads were subsequently aligned to the GRCh38 human genome using STAR [45] v2.7.7a with the following parameters: *--outSAMstrandField intronMotif --outSAMattrIHstart 0 --outFilterType BySJout --outFilterMultimapNmax 100 --winAnchorMultimapNmax 100 --alignSJoverhangMin 8 -- alignSJDBoverhangMin 1 --outFilterMismatchNmax 999 –outFilterMismatchNoverLmax 0.04 --alignIntronMin 20 --alignIntronMax 1000000 --alignMatesGapMax 1000000 --outFilterIntronMotifs RemoveNoncanonical --readFilesCommand zcat --peOverlapNbasesMin 50*. Because pull-down–derived RNA libraries typically undergo additional PCR amplification, PCR duplicates were removed using Picard MarkDuplicates v2.24.1 (https://broadinstitute.github.io/picard/) and resulted BAM files were further filtered to retain only alignments from properly paired reads using samtools v1.7 [46] with *view -f 2* parameters. Then gene level quantifications were performed with HTSeq-count [47] v0.13.5 using Ensembl GTF gene annotation related to release 99 [48] and these parameters: *-s reverse -m union -t exon*. Read counts at each processing stage are summarized in the accompanying prospect (Supplementary Table 2).

Genes were filtered to retain those with at least 10 raw counts in at least 2 samples. Filtered counts were used to construct an edgeR v3.34.1 [49] DGEList object. Library-size normalization factors were computed using calcNormFactors(“none”) imposing no between-sample normalization. Normalized expression matrices were generated as CPM using edgeR::cpm and edgeR::rpkm. Gene lengths used to calculate FPKMs were computed from the Ensembl release 99 GTF as the total length of the union of all exonic intervals for each gene locus. Sample-to-sample similarity was assessed by computing a Pearson’s correlation matrix on CPM values, which was visualized using corrplot with hierarchical clustering order. For differential expression testing, a generalized linear model (GLM) framework was applied using edgeR. A design matrix was built using the following design *(∼0 + condition*). Dispersion parameters were estimated with estimateDisp function with robust=TRUE parameter to reduce the influence of potential outliers. The negative binomial GLM was fitted with glmFit, and differential expression was assessed using likelihood ratio tests (glmLRT) for: the contrasts: EVEN vs INP; ODD vs INP or LACZ vs INP. In each condition the candidates lncRNA interactors were defined as the RNAs enriched in both EVEN and ODD pulldown sets and not enriched in *LACZ* control.

Isoform quantification in iPSC-derived MN (GSE94888; [50]) and SK-N-BE cells (GSE244751; [23]) was performed using Salmon v1.10.3 [51].

### Statistical analyses

Data are presented as mean values, and error bars represent the standard error of the mean (SEM). The number of independent biological replicates (N) for each experiment is reported in the figure legends. Circles, triangles, squares and diamonds denote individual biological replicates. Errors were calculated from relative quantities (2^−ΔCt^) or normalized values (2^−ΔΔCt^) and appropriately propagated. Statistical significance was assessed using two-tailed paired, unpaired Student’s *t*-tests (performed in Microsoft Excel) or two-way ANOVA followed by Tukey’s post hoc test (α=0.05, performed in GraphPad Prism), as indicated in the figure legends. A *P*-value < 0.05 was considered statistically significant (\**P* < 0.05, \*\**P* < 0.01, \*\*\**P* < 0.001, *\*\*\*\*P* < 0.0001).

## RESULTS

### *LINC00205* is enriched in SG and interacts with FUS^P525L^

To identify lncRNAs involved in pathological SG assembly and dynamics in the molecular context of ALS, we employed SK-N-BE Neuroblastoma cells expressing the mutant FUS^P525L^ in a Doxycycline (DOX)-inducible manner and exposed them to oxidative stress through Sodium Arsenite (ARS) treatment for 1 hour. The P525L mutation disrupts the nuclear localization signal of FUS, leading to its cytoplasmic mislocalization and, upon stress conditions, to aberrant incorporation into SG with pathological solid-like properties [52–54]. We searched for lncRNAs enriched (log2 fold-enrichment (SGenr/INP) > 1 and FDR < 0.05) in FUS^P525L^-containing SG in the dataset produced by Mariani et al. [23]. Among the 1510 transcripts identified, we focused on the 139 classified as lncRNA (Fig. 1A, left panel). By cross-referencing this dataset with the results of a FUS^P525L^ HITS-CLIP experiment performed by Di Timoteo *et al.* [55] in the same cellular model, we identified 13 lncRNAs as FUS binders (Fig. 1A, right panel and Supplementary Table 3). Among these, *LINC00205* was chosen as the candidate of interest for further studies, based on four key features: i) its multi-exonic structure, a common characteristic among functionally relevant lncRNAs (Supplementary Table 3); ii) its relatively long sequence (7553 nucleotides), ranking among the three longest lncRNAs in the list, which provides a favorable premise for acting as a scaffold for RNAs and proteins during the nucleation and organization of SG structures [29,30] (Supplementary Table 3); iii) its relatively high levels of expression (average FPKM of 6,7 in FUS^P525L^ SK-N-BE) (Supplementary Table 3); and iv) its highest specificity for FUS^P525L^ binding compared with FUS^WT^ in human induced Pluripotent Stem Cells (hiPSC)-derived spinal Motor Neurons (MN) among the identified lncRNA candidates [50] (Supplementary Table 3).

**Fig. 1:**
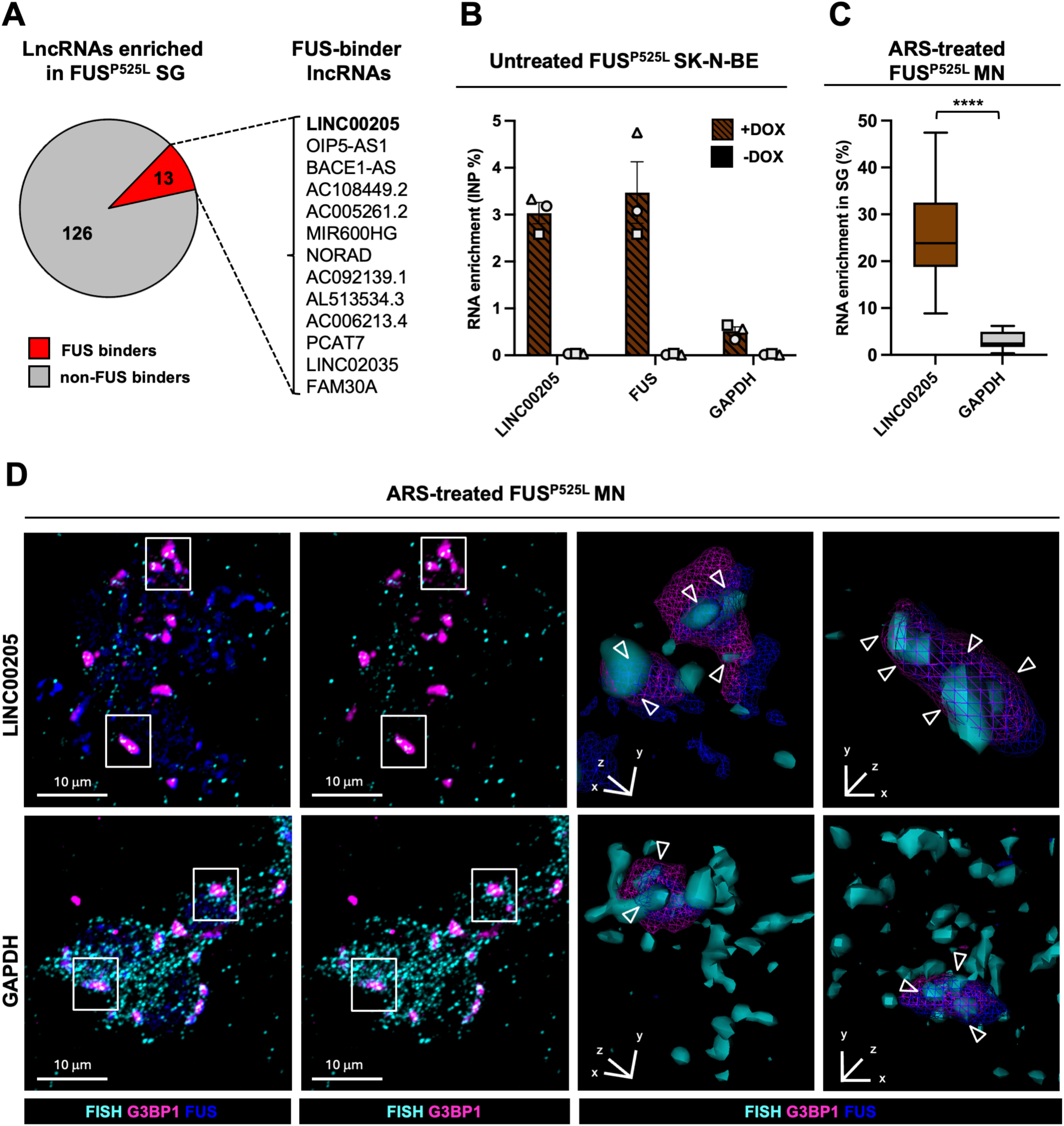
LINC00205 is enriched in SG and interacts with FUS^P525L^. **A)** Pie chart (left) showing the 139 lncRNAs enriched in FUS^P525L^ SG in SK-N-BE cells, categorized as FUS-binders and non-FUS binders. The FUS-binder lncRNAs are listed on the right*. LINC00205* is highlighted in bold. **B**) qRT-PCR analysis of *LINC00205* enrichment over INP from FLAG-FUS^P525L^ CLIP assay performed in untreated FUS^P525L^ SK-N-BE cells. -DOX cells were used as CLIP negative control. Data are expressed as mean of INP percentage ± SEM. *FUS* and *GAPDH* were used as positive and negative controls, respectively. N=3. **C)** Boxplot showing quantification of *LINC00205*/G3BP1-positive SG colocalization in ARS-treated FUS^P525L^ MN. *GAPDH* served as negative control. 242 and 227 cells were analyzed for *LINC00205* and *GAPDH* colocalization analysis, respectively. N=2. ****p≤0.0001 (two-tailed, unpaired Student’s T-test). **D)** Representative smFISH/immunofluorescence confocal images of ARS-treated FUS^P525L^ MN showing colocalization of *LINC00205* (upper-left panels) and *GAPDH* (lower-left panels) with G3BP1-positive SG. FUS staining marks nuclei. 3D renderings (upper and lower-right panels) are derived from indicated regions.

Interrogation of the Ensembl database (Archive Release 109, February 2023) revealed that the *LINC00205* gene is located on the long arm of chromosome 21 (21q22.3) and is annotated with the GENCODE gene identifier ENSG00000223768. *LINC00205* is transcribed from the forward strand and gives rise to five annotated splice variants (Supplementary Fig. 1A). Importantly, in iPSC-derived MN and ARS-treated SK-N-BE cells [23,50], the *ENST00000400362* isoform emerged as the most abundantly expressed variant (Supplementary Fig. 1B). This isoform represents the longest transcript variant (7553 nucleotides) and consists of three exons. All subsequent experiments were conducted to investigate the function of this specific *LINC00205* isoform. To validate the enrichment of *LINC00205* in pathological SG, we performed qRT-PCR analysis on RNAs purified from SG derived from FUS^P525L^ and FUS^WT^ SK-N-BE cells stressed with ARS [23]. Untreated cells were used as a technical control. *HUWE1*, a previously reported SG-enriched transcript [23,56], served as a positive control, whereas *ATP5O*, which is not associated with SG [23] was used as a negative control. Consistent with the RNA-seq data, *LINC00205* was significantly enriched in SG isolated from FUS^P525L^-expressing cells (Supplementary Fig. 1C). Notably, we found that *LINC00205* was also enriched in FUS^WT^ SG, indicating that it could play an important role in granule dynamics also in physiological conditions (Supplementary Fig. 1D).

To delineate a functional crosstalk between *LINC00205* and FUS, we tested their physical interaction performing a CLIP assay in SK-N-BE cells expressing FLAG-FUS^P525L^. Cells cultured in the absence of DOX (-DOX) were used as a negative control. The efficiency of FLAG-FUS^P525L^ CLIP was verified by WB (Supplementary Fig. 1E). The enrichment of *LINC00205* in the immunoprecipitation fractions over the input (INP) was evaluated by qRT-PCR, using *FUS* and *GAPDH* mRNAs as positive and negative controls, respectively (Fig. 1B). As expected, *LINC00205* was enriched at levels comparable to the positive control, confirming its interaction with FUS^P525L^ in the cytoplasm.

Even if SK-N-BE cells represent a useful model for investigating neuronal differentiation and neurodegenerative disorders, we next introduced spinal MN derived from hiPSC [38] as a more reliable cellular model system for ALS studies. Two different hiPSC lines were used: one expressing FUS^WT^ and another genetically engineered to endogenously express the mutant FUS^P525L^, which displays cytoplasmic delocalization [57]. *LINC00205* expression levels progressively increased during MN differentiation of FUS^WT^ and FUS^P525L^ cell lines (Supplementary Fig. 1F, left and right panels, respectively), reaching their highest levels in mature MN compared to undifferentiated stem cells. This expression pattern supported a functional role for *LINC00205* in MN biology and validated the use of iPSC-derived MN as a relevant cellular model to investigate its role in ALS.

SmFISH combined with immunofluorescence was used to assess the co-localization of *LINC00205* with MN SG marked by G3BP1, a core SG protein [58]. *LINC00205* was enriched within SG in both FUS^P525L^ (Fig. 1C, 1D and Supplementary Fig. 1G) and FUS^WT^ (Supplementary Fig. 1H, I and J) ARS-treated MN; this result is consistent with the enrichment of *LINC00205* detected in SG purified from SK-N-BE cells (Supplementary Fig. 1C and D). This enrichment was significantly higher than that detected for *GAPDH*, which was used as negative control (Fig. 1C, 1D and Supplementary Fig. 1G; Supplementary Fig. 1H, I and J).

### *LINC00205* depletion reduces the number of FUS^P525L^-containing SG in human MN

To investigate the involvement of *LINC00205* in modulating pathological SG dynamics, we implemented a loss-of-function approach. Specifically, a *LINC00205* knock-out (KO) was generated using a CRISPR/Cas9-based strategy. A cassette containing a Poly-adenylation sequence (PAS) and a Neomycin resistance gene was inserted downstream of exon 1 of the *LINC00205* locus in both FUS^WT^ and FUS^P525L^ hiPSC lines (Supplementary Fig. 2A), resulting in the premature termination of *LINC00205* transcription. Two independent homozygous KO clones showing complete downregulation of the lncRNA were obtained for both FUS^WT^ (wt-KO#1 and wt-KO#2) and FUS^P525L^ (525-KO#1 and 525-KO#2) cell lines (Supplementary Fig. 2B).

To confirm that the depletion of *LINC00205* did not alter the differentiation process of hiPSC into MN, we assessed the expression of three differentiation markers by qRT-PCR: OCT4, a key regulator of stem cell pluripotency [59]; HB9, a MN progenitor marker [60]; and ISL1, involved in the specification of MN [61]. Comparison between CTRL and *LINC00205* KO conditions in both FUS^WT^ (Supplementary Fig. 2C) and FUS^P525L^ (Supplementary Fig. 2D) MN across multiple stages of differentiation demonstrated that *LINC00205* depletion did not alter the differentiation process.

To investigate the role of *LINC00205* in SG dynamics and to determine whether its function differs between physiological and pathological conditions, we performed immunofluorescence staining for G3BP1 and FUS in ARS-treated FUS^P525L^ and FUS^WT^ MN to visualize and quantify the number of SG per cell in CTRL and *LINC00205* KO conditions. In FUS^P525L^ MN, depletion of *LINC00205* led to a significant reduction (∼30%) in the total number of G3BP1-containing SG (total G3BP1 SG) per cell (525-KO#1 and 525-KO#2), compared to 525-CTRL (Fig. 2A, left panel). A comparable decrease was observed when quantifying the total number of FUS^P525L^-marked SG (total FUS^P525L^ SG) per cell under the same conditions (Fig. 2A, right panel). To determine whether this effect specifically involved FUS^P525L^-containing SG, we classified all G3BP1 SG into two populations: i) G3BP1/FUS^P525L^ SG, positive for both G3BP1 and FUS^P525L^, and ii) G3BP1-only SG, which do not contain FUS^P525L^. Interestingly, in FUS^P525L^ MN the number of G3BP1/FUS^P525L^ SG per cell was significantly reduced upon KO of *LINC00205* compared to 525-CTRL (Fig. 2B, left panel), whereas the number of G3BP1-only SG was not affected (Fig. 2B, right panel). The effect is further described by representative images shown in Fig. 2C. In contrast, in FUS^WT^ MN, where FUS in maintained in the nucleus and does not migrate into SG, the number of total G3BP1 SG was not altered by *LINC00205* depletion (Fig. 2D and S2E). These findings indicate that *LINC00205* specifically contributes to the formation of FUS^P525L^-containing SG without influencing the number of G3BP1-only SG.

**Fig. 2:**
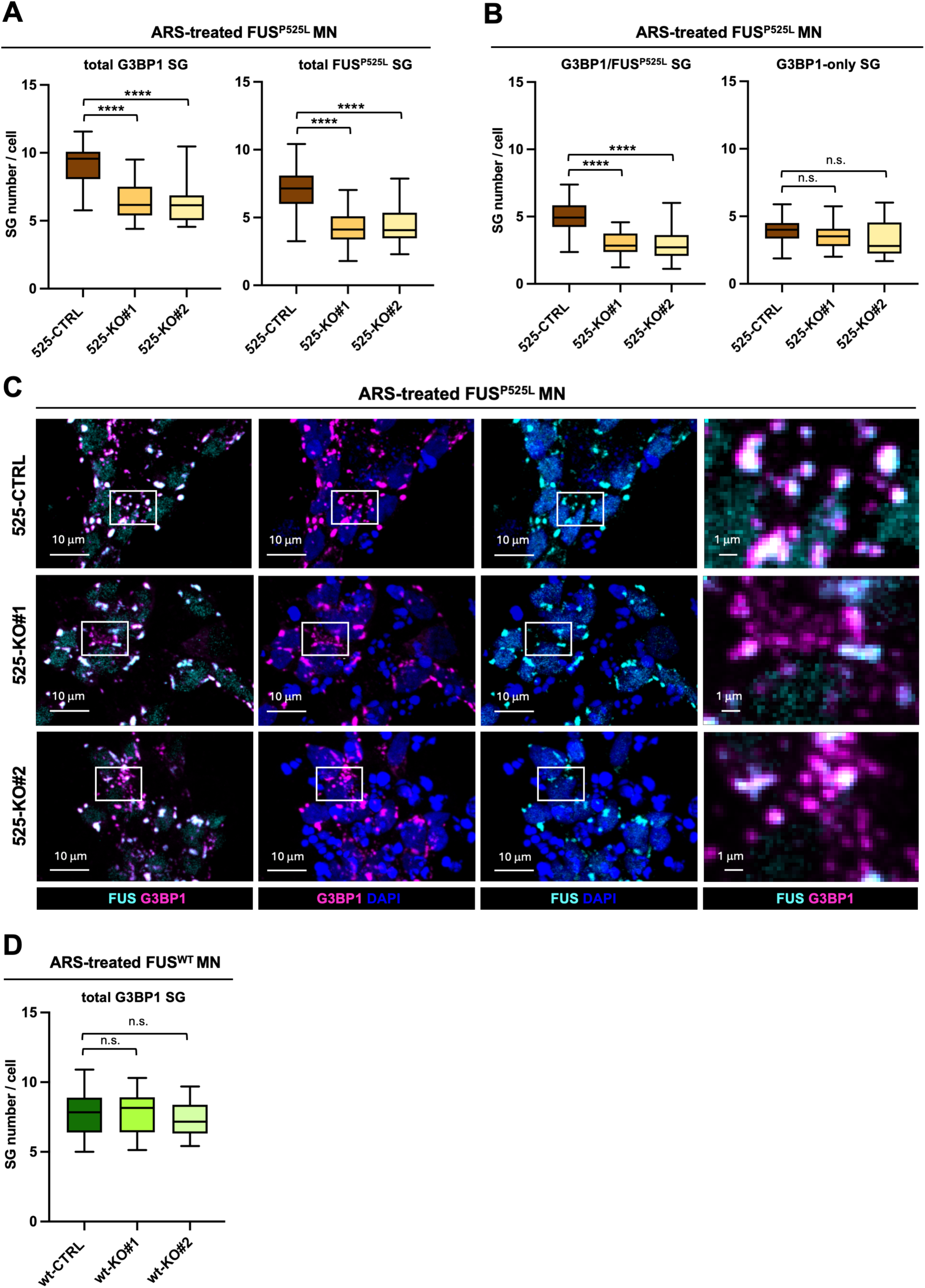
*LINC00205* depletion reduces the number of FUS^P525L^-containing SG in MN. **A)** Boxplot showing quantification of the total number of G3BP1-positive SG per cell (left panel) and FUS^P525L^-positive SG per cell (right panel) in ARS-treated 525-KO#1 and 525-KO#2 MN compared to 525-CTRL. 519, 687 and 319 cells were analyzed for 525-CTRL, 525-KO#1 and 525-KO#2 MN, respectively. N=3. ****p≤0.0001 (two-tailed, unpaired Student’s T-test). **B)** Boxplot showing quantification of the total number of G3BP1/FUS^P525L^-positive SG per cell (left panel) and G3BP1-only SG per cell (right) in ARS-treated 525-KO#1 and 525-KO#2 MN compared to 525-CTRL. Details as in A. ****p≤0.0001; n.s. p>0.05 (two-tailed, unpaired Student’s T-test). **C)** Representative immunofluorescence confocal images of ARS-treated FUS^P525L^ MN showing SG marked by both G3BP1 and FUS^P525L^ in 525-CTRL (upper row), 525-KO#1 (middle row) and 525-KO#2 (lower row) cells. Nuclei were counterstained with DAPI. Right column panels show magnifications of the indicated regions. **D)** Boxplot showing quantification of the total number of G3BP1-positive SG per cell in ARS-treated wt-KO#1 and wt-KO#2 MN compared to wt-CTRL. 283, 497 and 370 cells were analyzed for wt-CTRL, wt-KO#1 and wt-KO#2 MN, respectively. N=3. n.s. p>0.05 (two-tailed, unpaired Student’s T-test).

To determine whether *LINC00205* regulates FUS^P525L^ aggregation into SG without affecting FUS^P525L^ expression, we measured FUS RNA and protein levels in FUS^P525L^ MN. We found that neither RNA nor protein levels were significantly altered following *LINC00205* KO compared to control conditions, both before and after stress (Supplementary Fig. 2F and G, left and right panels). As an additional control, no changes in FUS RNA or protein levels were observed in FUS^WT^ MN (Supplementary Fig. 2H and I, left and right panels). These results indicate that *LINC00205* does not regulate FUS expression but rather acts downstream of protein synthesis to promote the formation of FUS^P525L^ -containing SG.

### *LINC00205* loss favors faster SG recovery in FUS^P525L^ MN

Given that mutant FUS enrichment within SG is known to enhance their solidification [54,62] we hypothesized that the reduction of FUS^P525L^-containing SG upon *LINC00205* KO could also affect SG recovery. To test this hypothesis, MN expressing either FUS^P525L^ or FUS^WT^, under CTRL and *LINC00205*-depleted conditions, were analyzed at multiple time points.

Cells were treated with ARS for 1 hour and subsequently examined after 1, 2, and 4 hours of recovery following ARS removal (1, 2, and 4h REC). Using G3BP1 immunofluorescence, we quantified the percentage of MN retaining at least one SG to assess SG recovery kinetics. As expected, FUS^P525L^ MN remained nearly 100% granulated at all recovery time points (525-CTRL), whereas 525-KO#1 and 525-KO#2 cells exhibited a progressive reduction in the proportion of granulated cells, with decreases of approximately 30% and 40% after 2 and 4 hours of recovery, respectively (Fig. 3A and C). As control, wt-KO#1 and wt-KO#2 displayed recovery kinetics comparable to wt-CTRL MN (Fig. 3B and Supplementary Fig. 3A).

**Fig. 3:**
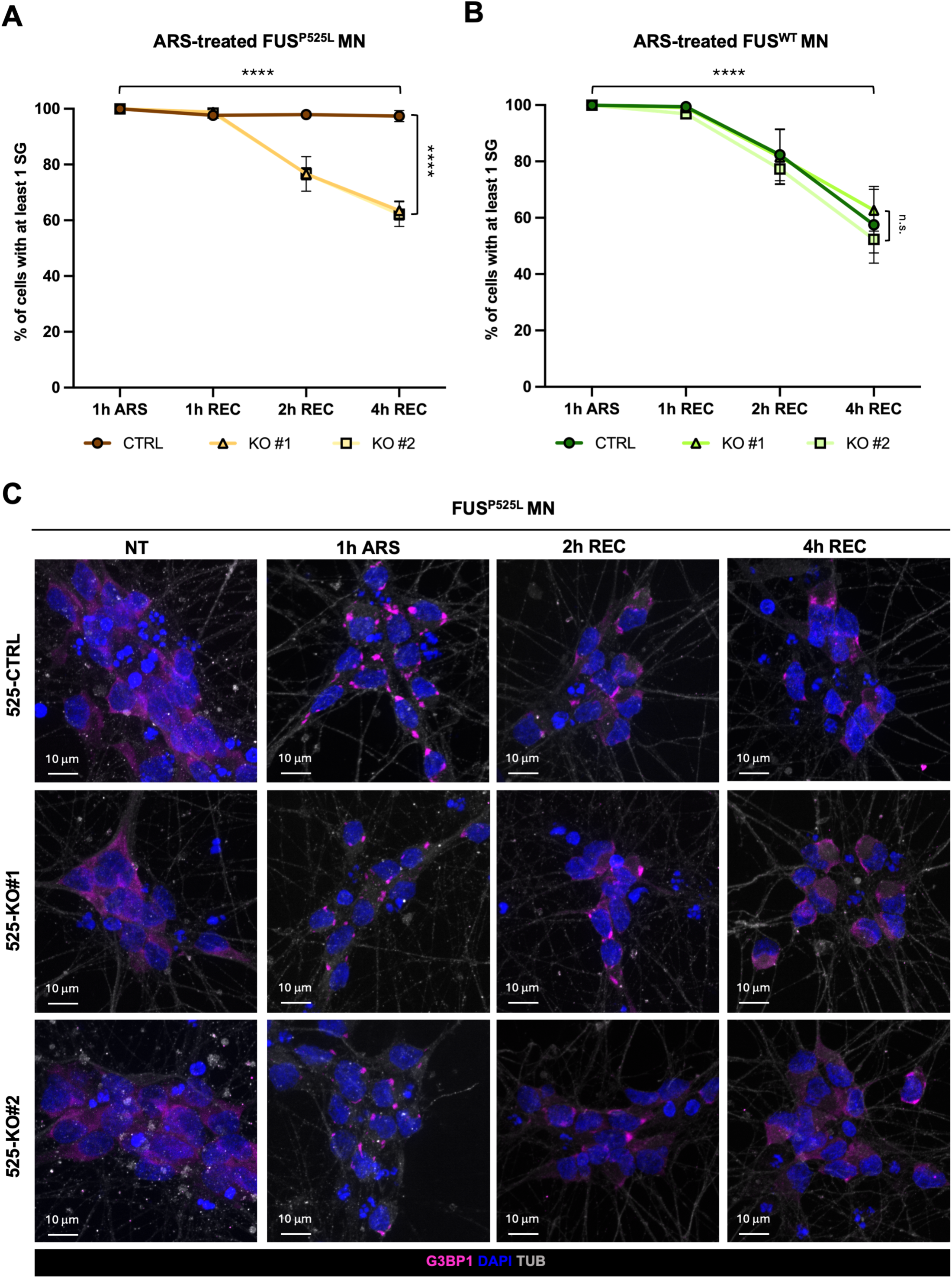
*LINC00205* loss favors faster SG recovery in FUS^P525L^ MN. **A)** Line graph showing the percentage of ARS-treated FUS^P525L^ MN containing at least one G3BP1-positive SG at different recovery (REC) time points. The numbers of cells analyzed for 1h ARS, 1, 2 and 4h REC, respectively, were: 171, 483, 553, and 491 for 525-CTRL; 507, 759, 468, and 355 for 525-KO#1; 404, 410, 375, and 440 for 525-KO#2. Data are expressed as mean ± SEM. N=3. ****p≤0.0001 (two-way ANOVA test). Statistical significance is summarized in Supplementary Table 4. **B)** Line graph showing the percentage of ARS-treated FUS^WT^ MN containing at least one G3BP1-positive SG at different recovery (REC) time-points. The numbers of cells analyzed for 1h ARS, 1, 2 and 4h REC, respectively, were: 398, 544, 365, and 735 for wt-CTRL; 184, 171, 204, and 428 for wt-KO#1; 267, 468, 393, and 509 for wt-KO#2, respectively. Data are expressed as mean ± SEM. N=3. **** p≤0.0001, n.s. p>0.05 (two-way ANOVA test). Statistical significance is summarized in Supplementary Table 4. **C)** Representative immunofluorescence confocal images of FUS^P525L^ MN at different time points (NT, 1h ARS, 2h REC, 4h REC, each column) in 525-CTRL (upper row), 525-KO#1 (middle row) and 525-KO#2 (lower row) cells. Nuclei were counterstained with DAPI.

Notably, after 4 hours of recovery, the percentage of recovered 525-KO#1 and 525-KO#2 MN was comparable to that observed under FUS^WT^ conditions, demonstrating that *LINC00205* depletion in FUS^P525L^ cells restores a physiological rate of SG dissolution. Overall, these data indicate an important role of *LINC00205* in regulating SG dynamics specifically under pathological conditions.

### *LINC00205* promotes selective mRNA localization to FUS^P525L^-containing SG

LncRNAs are increasingly recognized as key molecular scaffolds that provide structural frameworks for the coordinated assembly of proteins and RNAs, thereby facilitating the dynamic formation and regulation of membrane-less organelles in response to cellular stress or environmental changes [63–65]. Having established that *LINC00205* directly interacts with FUS^P525L^ and promotes the condensation of FUS^P525L^-containing SG, we sought to identify additional components of the macromolecular complex controlled by *LINC00205* in the formation of pathological SG.

To this end, we first identified *LINC00205*-associated RNAs using a native RNA Pull-Down (PD) assay followed by RNA-seq in SK-N-BE cells expressing either FUS^WT^ or FUS^P525L^, in untreated and ARS-treated conditions. PD efficiency was quite high (10-20%) as assessed by qRT-PCR of *LINC00205* (Supplementary Fig. 4A and B). The PD RNA-seq analysis revealed that the lncRNA exhibits a different set of interactors depending on the FUS variant (WT *versus* P525L) and the cellular stress condition (untreated *versus* ARS-treated) (Supplementary Fig. 4C). Interestingly, in ARS-treated FUS^P525L^ cells, 91 RNAs were found to specifically associate with *LINC00205*, a substantially higher number compared with the 51 RNAs distinctively present in the PD performed in FUS^WT^ stressed cells (Supplementary Fig. 4D). These results suggest that the lncRNA engages with an increased number of RNA partners when FUS^P525L^ is present.

Among the 91 RNAs binding to *LINC00205* exclusively in FUS^P525L^cells, we selected 18 RNAs previously reported by Mariani and Setti [23] to be specifically enriched in FUS^P525L^-containing SG (Fig. 4A and Supplementary Table 5). Two of these were found to bind *LINC00205* already before stress (group 1, Supplementary Table 5), while 16 interacted with *LINC00205* exclusively upon stress (group 2, Supplementary Table 5).

**Fig. 4:**
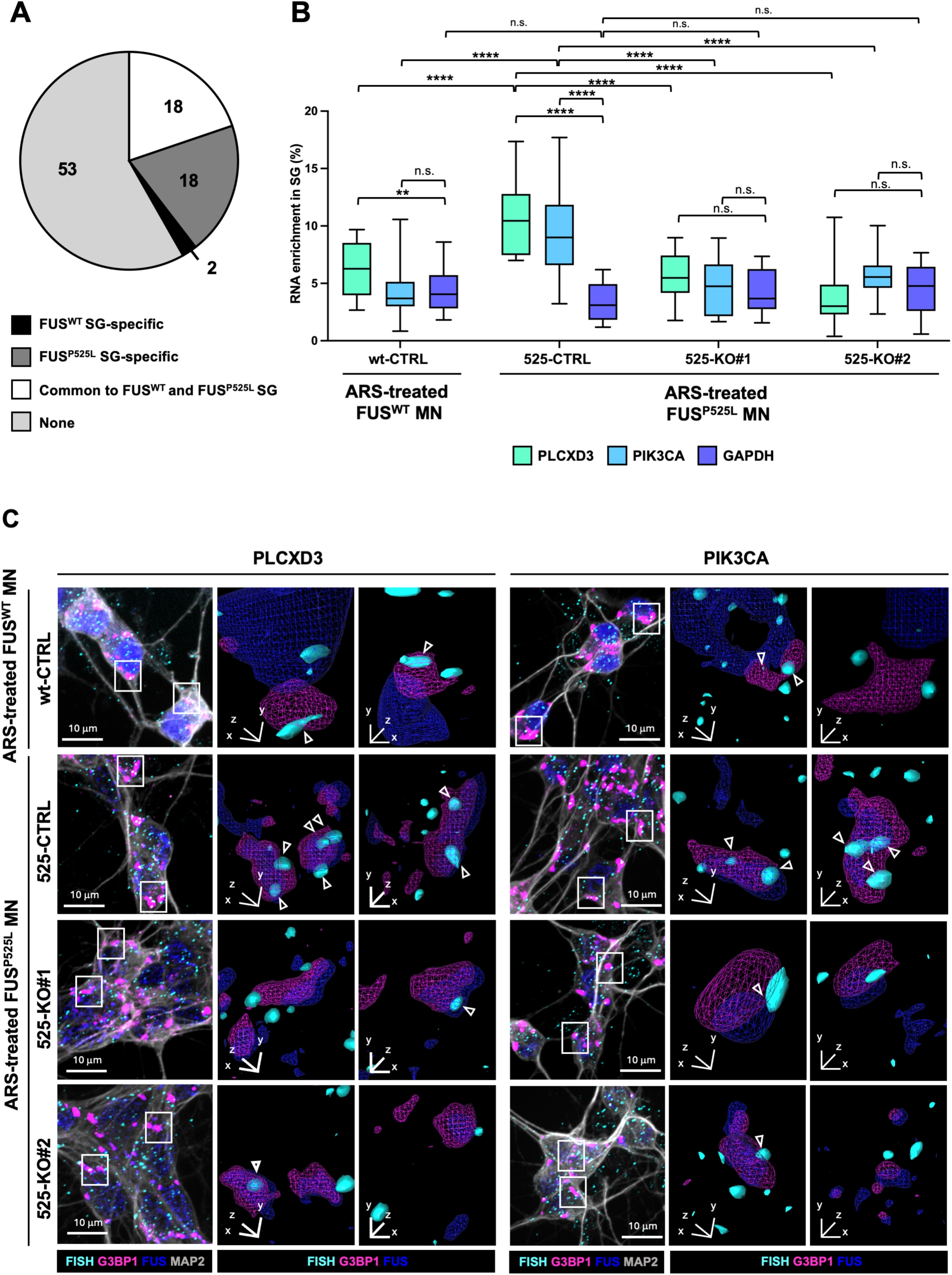
*LINC00205* promotes selective mRNA localization to FUS^P525L^-containing SG. **A)** Pie chart illustrating the distribution of the 91 RNAs specifically bound to *LINC00205* in ARS-treated FUS^P525L^ SK-N-BE cells, with respect to RNAs enriched in FUS^WT^ and FUS^P525L^ SG. **B)** Boxplot showing *PLCXD3* and *PIK3CA* enrichment in G3BP1-positive SG in FUS^WT^ (wt-CTRL) and FUS^P525L^ (525-CTRL, 525-KO#1 and 525-KO#2) MN. The numbers of cells analyzed for wt-CTRL, 525-CTRL, 525-KO#1, 525-KO#2, respectively, were: 156, 299, 320, and 232 for *PLCXD3*; 218, 231, 227, and 236 for *PIK3CA*; 193, 227, 206, and 214 for *GAPDH* (N = 2). **p≤0.01, ****p≤0.0001; n.s. p>0.05 (two-tailed, unpaired Student’s T-test). **C)** Representative smFISH/immunofluorescence confocal images of ARS-treated FUS^WT^ and FUS^P525L^ MN showing colocalization of *PLCXD3* (left panels) and *PIK3CA* (right panels) with G3BP1-positive SG in wt-CTRL (upper row) 525-CTRL (middle-upper row), 525-KO#1 (middle-lower row) and 525-KO#2 (lower row) cells. FUS staining marks nuclei. 3D renderings (middle-right and right columns) are derived from indicated regions.

*PLCXD3* [66] and *ZNF841* (Razavi,R., Fathi,A., Yellan,I., *et al.* Extensive binding of uncharacterized human transcription factors to genomic dark matter. *bioRxiv*, 2024, 10.1101/2024.11.11.622123), belonging to group 1, and *PIK3CA* [67] and *RAB30* [68], belonging to group 2, were selected as candidates for validation in independent RNA pull-down experiments in FUS^P525L^ and FUS^WT^ cells before and after ARS treatment. The results show that all these mRNAs specifically interact with *LINC00205* exclusively in FUS^P525L^ conditions, both before and after stress (Supplementary Fig. 4E, F, G and H).

We selected for further analysis *PLCXD3*, a member of the phosphoinositide-specific phospholipases family 6 [66,69], and *PIK3CA*, which encodes a major regulator of cellular redox homeostasis [70]. We proceeded to evaluate the effects of *LINC00205* on the SG localization of these transcripts by performing smFISH combined with immunofluorescence for G3BP1 and FUS, in MN expressing either FUS^P525L^ or FUS^WT^. *PLCXD3* and *PIK3CA* were significantly enriched in FUS^P525L^ SG (∼11% and ∼10%, respectively) compared to FUS^WT^ SG (∼6% and ∼4%, respectively) and relative to *GAPDH* mRNA, used as a negative control (Fig. 4B and C and Supplementary Fig. 4I). Notably, upon *LINC00205* depletion, *PLCXD3* colocalization with SG was significantly reduced from 11% to 6% in 525-KO#1 and to 4% in 525-KO#2 MN (Fig. 4B and C). Similarly, *PIK3CA* colocalization with SG decreased from 10% to 5% in 525-KO#1 and to 6% in 525-KO#2 MN (Fig. 4B and C). Interestingly, the percentages of localization identified in 525-KO cells were comparable to those observed in FUS^WT^ conditions and to the *GAPDH* control (Fig. 4B and C and Supplementary Fig. 4I). These data indicate that *LINC00205* promotes the localization of its interactor mRNAs *PLCXD3* and *PIK3CA,* in FUS^P525L^-containing SG.

### *LINC00205* directly interacts with DHX36 and increases its colocalization with FUS^P525L^-containing SG

We next aimed to characterize the proteins participating in the macromolecular complex orchestrated by *LINC00205* during the formation of pathological SG. Based on previous findings by Mariani and Setti [23], we focused on DHX36 and HuR, that are enriched in FUS^P525L^-containing SG. DHX36 is a DEAH/RHA helicase [71,72], whereas HuR (ELAVL1) is an RNA-binding protein that binds AU-rich elements in mRNAs, thereby stabilizing them and modulating their translation [73–75]. We first assessed whether DHX36 and HuR directly bind *LINC00205* by performing CLIP assays in untreated SK-N-BE cells expressing FUS^P525L^. The efficiency of DHX36 (Supplementary Fig. 5A) and HuR (Supplementary Fig. 5B, left panel) immunoprecipitation (IP) was verified by WB. The enrichment of *LINC00205* in the IP fractions relative to the INP was assessed by qRT-PCR, using *WBP4* and *RPS7* mRNAs as positive and negative controls, respectively, for DHX36 (Fig. 5A), and *HUR* and *GAPDH* mRNAs as positive and negative controls, respectively, for HuR (Supplementary Fig. 5B, right panel). The results indicate that *LINC00205* is enriched in the DHX36 IP fractions (Fig. 5A); in contrast, no association between *LINC00205* and HuR was detected (Supplementary Fig. 5B, right panel).

**Fig. 5:**
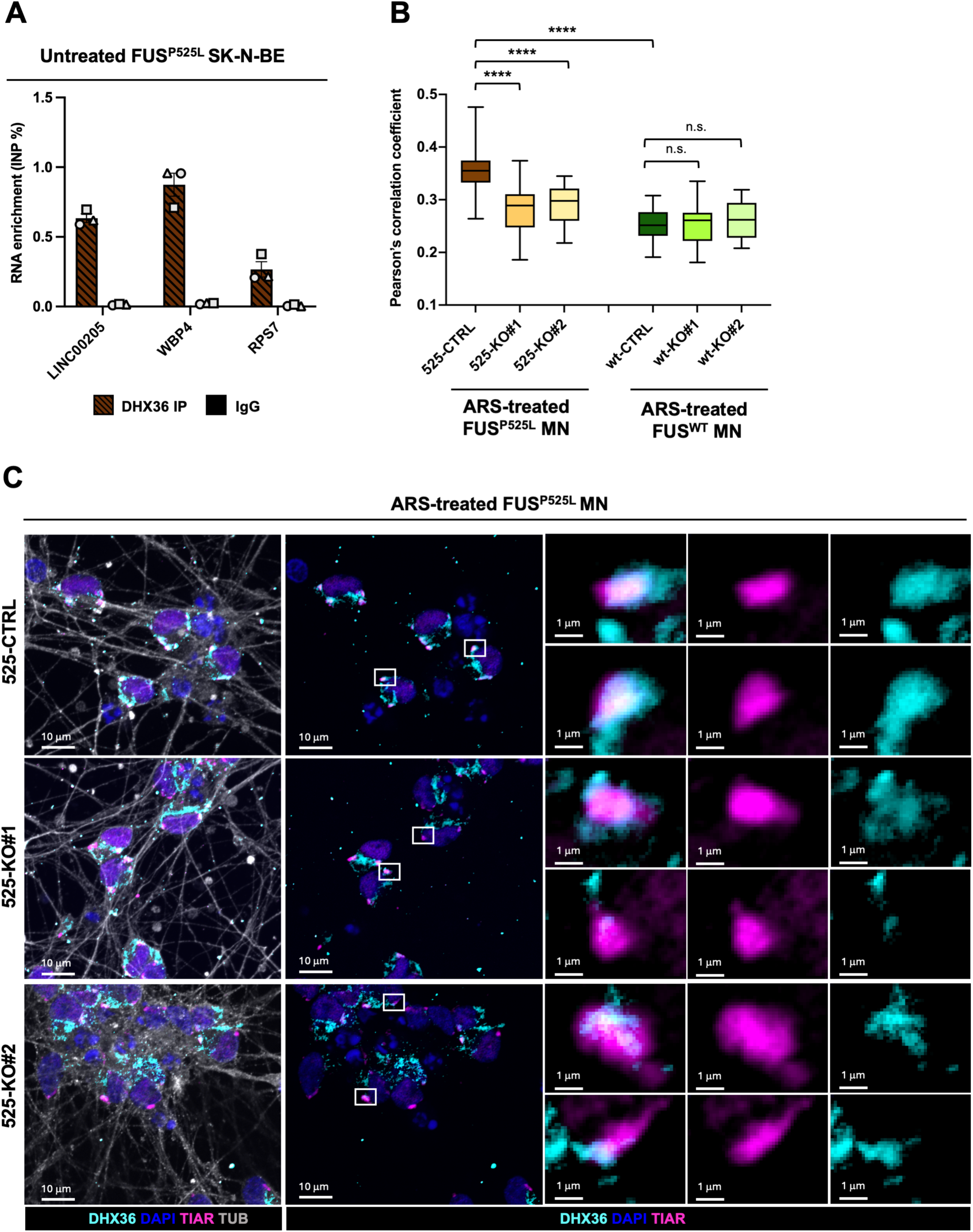
*LINC00205* directly interacts with DHX36 and increases its colocalization with FUS^P525L^-containing SG. **A)** qRT-PCR analysis of *LINC00205* enrichment over INP in IP and IgG fractions from DHX36 CLIP assay performed in untreated FUS^P525L^ SK-N-BE cells. Data are expressed as mean of INP percentage ± SEM. *WBP4* and *RPS7* were used as positive and negative controls, respectively. N=3. **B)** Boxplot showing Pearson’s correlation coefficients indicating DHX36/TIAR-positive SG colocalization by immunofluorescence analyses in ARS-treated FUS^P525L^ and FUS^WT^ MN. For FUS^P525L^ and FUS^WT^ MN conditions (CTRL, KO#1, KO#2), the number of cells analyzed were 407, 365, 513, and 282, 202, 259, respectively. N=2. ****p≤0.0001; n.s. p>0.05 (two-tailed, unpaired Student’s T-test). **C)** Representative immunofluorescence confocal images of ARS-treated FUS^P525L^ MN showing DHX36/TIAR-positive SG colocalization in 525-CTRL (upper row), 525-KO#1 (middle row) and 525-KO#2 (lower row) cells. Nuclei were counterstained with DAPI. Right panels show magnifications of the indicated regions. All images represent a single focal plane from the acquired z-stacks.

To investigate whether *LINC00205* influences DHX36 colocalization with pathological SG, we performed immunofluorescence staining for DHX36 and TIAR, a SG marker [76,77], in ARS-treated FUS^P525L^ and FUS^WT^ MN in CTRL and *LINC00205* KO conditions. Consistent with previous reports by Mariani and Setti [23], DHX36 was significantly more enriched in FUS^P525L^-containing SG compared to FUS^WT^ SG (Fig. 5B, 5C and Supplementary Fig. 5C). Notably, depletion of *LINC00205* reduced DHX36 levels in FUS^P525L^ SG to amounts comparable to those observed in FUS^WT^ SG (Fig. 5B). As control, DHX36 enrichment in FUS^WT^ SG was unaffected by *LINC00205* KO relative to the CTRL condition (Fig. 5B). We also assessed HuR colocalization within SG marked by G3BP1 in the same conditions. In line with the lack of direct binding between *LINC00205* and HuR, we observed that HuR abundance in FUS^P525L^ and FUS^WT^ SG remained unchanged upon *LINC00205* depletion (Supplementary Fig. 5D, E and F). Taken together, these findings suggest that *LINC00205* directly binds the DHX36 protein and enhances its localization to pathological SG, whereas it has no effect on the SG localization of HuR, which does not associate with the lncRNA.

## DISCUSSION

In this study, we identify *LINC00205* as a regulator of pathological SG composition and dynamics in the context of the ALS-linked FUS^P525L^ mutation.

By integrating RNA-seq analysis of purified SG, CLIP and RNA pull-down, loss-of-function and imaging approaches, we uncovered a mechanistic role for *LINC00205* as an RNA scaffold that selectively promotes the assembly and persistence of FUS^P525L^-containing SG, thereby modulating their composition and disassembly kinetics.

A first key finding of our work is that *LINC00205*, which we demonstrated to directly interact with FUS^P525L^, is robustly enriched in SG, both in neuronal cells and in human iPSC-derived MN. Although its localization to SG occurs independently of mutant FUS, our data reveal that *LINC00205* plays a distinct functional role specifically in the context of FUS^P525L^-containing pathological assemblies.

The specificity of *LINC00205* function becomes particularly evident upon its genetic ablation. *LINC00205* depletion selectively reduces the number of FUS^P525L^ SG without affecting the formation of canonical G3BP1 SG or altering FUS expression levels. This indicates that *LINC00205* acts downstream of mutant FUS mislocalization and does not broadly impair the cellular stress response. Instead, *LINC00205* appears to control SG RNA and protein composition, specifically favoring the intake of FUS^P525L^ in SG. This distinction is particularly relevant considering increasing evidence that not all SG are equivalent, and that disease-associated RBP mutations confer aberrant physical properties to specific SG subpopulations [20,57,78,79].

Consistent with this notion, loss of *LINC00205* markedly accelerates SG disassembly during stress recovery in FUS^P525L^ MN, restoring kinetics comparable to those observed in FUS^WT^ cells. Given that mutant FUS is known to promote a transition toward more solid-like, persistent SG, our findings suggest that *LINC00205* contributes to this pathological process. By reducing the fraction of mutant FUS-containing SG, *LINC00205* depletion may facilitate granule dissolution, thereby promoting a return to physiological cytoplasmic homeostasis.

At the molecular level, integrative analysis of RNA-seq data from *LINC00205* RNA PD and SG purification reveals that *LINC00205* preferentially associates with transcripts enriched in FUS^P525L^-containing SG. Among these, *PLCXD3* and *PIK3CA* mRNAs serve as examples *LINC00205* interactors, both prior to and after stress. Notably, the observation that *LINC00205* promotes *PLCXD3* and *PIK3CA* sequestration into pathological SG supports a model in which the lncRNA contributes to translational repression of selected mRNAs during stress, with potential consequences for cellular homeostasis.

In addition to RNA partners, we identify DHX36 as a protein component of the *LINC00205*-dependent SG network. Although both DHX36, a helicase involved in RNA stability, translation and phase separation [71,72,80], and HuR, an RNA-binding protein that regulates mRNA metabolism during cellular stress [81,82] are enriched in SG, with a higher propensity for FUS^P525L^-containing assemblies [23], only DHX36 directly interacts with *LINC00205*. Strikingly, *LINC00205* selectively promotes DHX36 accumulation within FUS^P525L^-containing pathological SG, while leaving HuR localization unaffected. This selective recruitment suggests that *LINC00205* may promote the enrichment of specific classes of RNAs within pathological SG. Notably, DHX36 localization to SG is restored to physiological levels upon *LINC00205* depletion, further supporting a scaffold function for the lncRNA in organizing SG composition rather than globally altering protein abundance.

Taken together, our findings support a model in which *LINC00205* acts as a multivalent RNA scaffold that contributes to the nucleation of pathological SG by simultaneously binding mutant FUS, selected mRNAs such as *PLCXD3* and *PIK3CA* and regulatory RBPs including DHX36. Through these interactions, *LINC00205* favors the selective condensation and persistence of mutant FUS-containing SG, contributing to their impaired dynamics characteristic of ALS-linked FUS mutations (Fig. 6A, left panel). Upon *LINC00205* depletion, this molecular platform is lost: mutant FUS-containing SG are markedly reduced, along with the recruitment of *PLCXD3* and DHX36 to SG. Therefore, SG display more physiological properties, including a recovery rate comparable to that observed in FUS^WT^-expressing cells (Fig. 6A, right panel).

**Fig. 6:**
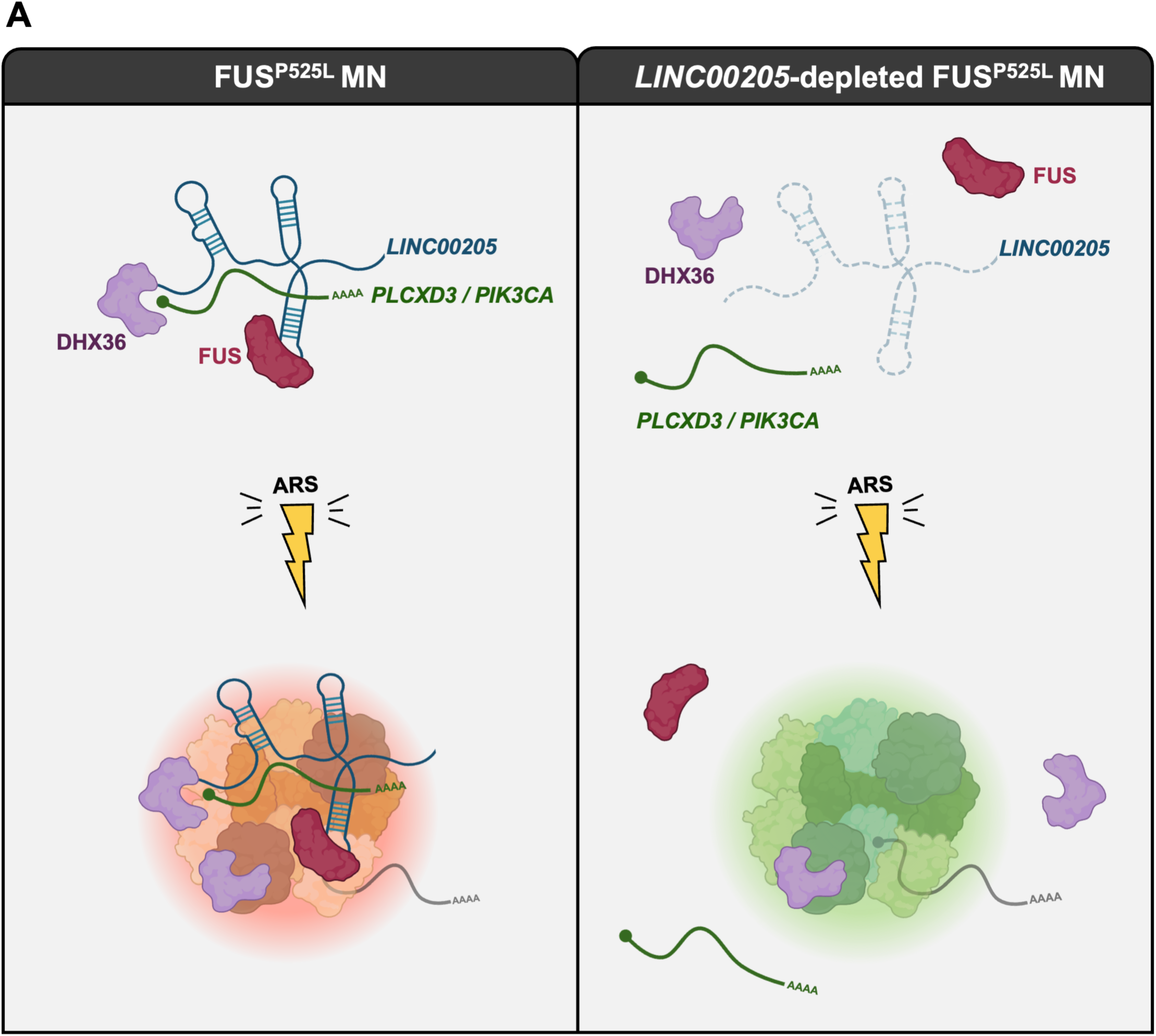
*LINC00205* mechanism of action in FUS^P525L^ MN. **A)** Right panel: in the cytoplasm, *LINC00205* interacts with FUS^P525L^ and scaffolds the assembly of a pathological ribonucleoprotein complex that also includes *PLCXD3* and *PIK3CA* mRNAs and DHX36. Upon cellular stress, this complex is recruited into SG, contributing to their impaired dynamics characteristics. Left panel: upon *LINC00205* depletion, the assembly of this pathological ribonucleoprotein complex is impaired: mutant FUS-containing SG are strongly reduced, together with the localization of *PLCXD3* and *PIK3CA* mRNAs and DHX36 to SG. Therefore, SG exhibit more physiological features, including a recovery kinetics like that observed in FUS^WT^-expressing cells.

While a limited number of lncRNAs, including *NEAT1* and *NORAD*, have been reported to influence cellular stress responses or to localize to SG [31,32,83–85], their roles appear largely indirect. Conversely, we have been able to show a precise function for *LINC00205* as a molecular scaffold that actively organizes the composition and persistence of ALS-linked pathological SG by driving specific RNA and protein components.

From a broader perspective, this work expands the functional repertoire of lncRNAs in neurodegeneration, highlighting their capacity to regulate phase-separated assemblies at multiple molecular levels. While our study focuses on FUS-associated ALS, similar RNA-mediated scaffolding mechanisms may operate in other proteinopathies characterized by aberrant RNP granules. Future work will be required to determine whether *LINC00205* modulation affects long-term neuronal survival and whether its activity is altered in patient-derived tissues.

Nonetheless, our data identify *LINC00205* as a critical determinant of pathological SG behavior and suggest that targeting lncRNA-dependent interaction networks may represent a novel strategy to restore proteostasis in ALS.

## ETHICS APPROVAL

Our research complies with all relevant ethical regulations.

Human neuroblastoma SK-N-BE cells were obtained from the American Type Culture Collection (ATCC). Human iPSC lines were generated as previously described in [57]. As reported in the original study, informed consent was obtained from all participants prior to cell donation.

## CONSENT FOR PUBLICATION

Not applicable.

## DATA AVAILABILITY

All software, links to websites or tools used for this work are referred to in the ‘Methods’ section. The RNA-Seq data presented in this study are available in GEO, accession number GSE319004. To review GEO accession number GSE319004, please visit and enter the token ghmhwkaarhwvpmx.

## AUTHORS’ CONTRIBUTIONS

JR: conceptualization, data curation, formal analysis, investigation, methodology, supervision, visualization, writing-original draft (main text) and revised manuscript. GS: data curation, formal analysis, investigation, methodology, visualization. TS: methodology, data curation, visualization. AS: bioinformatic analysis, formal analysis, data curation, software. MB and PC: investigation, methodology, writing-original draft (main text). LF: investigation, visualization. DM, PT and EV: methodology, data curation. IB: conceptualization, supervision, writing-original draft, funding acquisition, project administration. All authors approved the final version of the manuscript.

## ACKNOWLEDGEMENTS

We thank Dr. Pietro Laneve, Prof. Mariangela Morlando, and Dr. Manuel Beltran for insightful discussions and valuable suggestions. We are grateful to Marcella Marchioni and Dr. Alessandra Galati for technical assistance, and to Manuela Caruso for her support, as well as to the technical and administrative staff of the IIT for their organizational assistance.

We also acknowledge the Imaging Facility of CLN²S@Sapienza, Istituto Italiano di Tecnologia, Rome, and Dr. Valeria De Turris for support with confocal microscopy acquisitions. We acknowledge the Microscopy Facility of the Department of Biology and Biotechnology “C. Darwin” for assistance with image acquisition, analysis, and technical support (the facility is part of the Sapienza Research Infrastructure). We also thank Dr. Diego Vozzi and the Genomics Facility of IIT for their assistance with RNA sequencing experiment, and the members of the IIT Technologies Flagship.

Graphical abstract was partially created with BioRender, with permission available at: https://app.biorender.com/illustrations/69af0685740423be2a434486?slideId=745f3680-f5ec-403d-a2a1-f84957de727b.

This work is dedicated to the memory of Massimo Arceci, a kind and generous friend.

## AUTHORS’ INFORMATION

**Jessica Rea**: Center for Life Nano-& Neuro-Science, Fondazione Istituto Italiano di Tecnologia, Viale Regina Elena 291, 00161, Rome, Italy.

**Gaia Stortini**: Center for Life Nano-& Neuro-Science, Fondazione Istituto Italiano di Tecnologia, Viale Regina Elena 291, 00161, Rome, Italy; Department of Biology and Biotechnologies "C. Darwin", Sapienza University of Rome, Piazzale Aldo Moro 5, 00185, Rome, Italy.

**Tiziana Santini**: Department of Biology and Biotechnologies "C. Darwin", Sapienza University of Rome, Piazzale Aldo Moro 5, 00185, Rome, Italy.

**Adriano Setti:** Department of Biology and Biotechnologies "C. Darwin", Sapienza University of Rome, Piazzale Aldo Moro 5, 00185, Rome, Italy.

**Marta Bernardi**: Department of Biology and Biotechnologies "C. Darwin", Sapienza University of Rome, Piazzale Aldo Moro 5, 00185, Rome, Italy; Present Address: The James Black Centre, King’s College London, 125 Coldharbour Lane, London SE5 9NU, UK.

**Pierpaolo Cantisani**: Department of Biology and Biotechnologies "C. Darwin", Sapienza University of Rome, Piazzale Aldo Moro 5, 00185, Rome, Italy; Present Address: Department of Information Engineering and Mathematics, University of Siena, Via Roma 56, 53100, Siena, Italy.

**Letizia Fucci**: Department of Biology and Biotechnologies "C. Darwin", Sapienza University of Rome, Piazzale Aldo Moro 5, 00185, Rome, Italy.

**Davide Mariani**: Department of Biology and Biotechnologies "C. Darwin", Sapienza University of Rome, Piazzale Aldo Moro 5, 00185, Rome, Italy; Center for Human Technologies, Fondazione Istituto Italiano di Tecnologia, Via Enrico Melen 83, 16153, Genoa, Italy.

**Paolo Tollis**: Center for Life Nano-& Neuro-Science, Fondazione Istituto Italiano di Tecnologia, Viale Regina Elena 291, 00161, Rome, Italy.

**Erika Vitiello**: Center for Human Technologies, Istituto Italiano di Tecnologia, Via Enrico Melen 83, 16153, Genoa, Italy; Present Address: Institute of Pharmacy and Molecular Biotechnology (IPMB), Heidelberg University, 69120 Heidelberg, Germany.

**Irene Bozzoni**: Center for Life Nano-& Neuro-Science, Fondazione Istituto Italiano di Tecnologia, Viale Regina Elena 291, 00161, Rome, Italy; Department of Biology and Biotechnologies "C. Darwin", Sapienza University of Rome, Piazzale Aldo Moro 5, 00185, Rome, Italy.

## CORRESPONDING AUTHOR

Correspondence to Irene Bozzoni.

## REFERENCES

1. Gomes E, Shorter J. The molecular language of membraneless organelles. J Biol Chem. 2019;294:7115–27. 10.1074/jbc.TM118.001192

2. Hirose T, Ninomiya K, Nakagawa S, Yamazaki T. A guide to membraneless organelles and their various roles in gene regulation. Nat Rev Mol Cell Biol. Nature Publishing Group; 2023;24:288–304. 10.1038/s41580-022-00558-8

3. Li Y, Liu Y, Yu X-Y, Xu Y, Pan X, Sun Y, et al. Membraneless organelles in health and disease: exploring the molecular basis, physiological roles and pathological implications. Sig Transduct Target Ther. Nature Publishing Group; 2024;9:305. 10.1038/s41392-024-02013-w

4. Alberti S. Phase separation in biology. Curr Biol. 2017;27:R1097–102. 10.1016/j.cub.2017.08.069

5. Nesterov SV, Ilyinsky NS, Uversky VN. Liquid-liquid phase separation as a common organizing principle of intracellular space and biomembranes providing dynamic adaptive responses. Biochim Biophys Acta Mol Cell Res. 2021;1868:119102. 10.1016/j.bbamcr.2021.119102

6. Boeynaems S, Alberti S, Fawzi NL, Mittag T, Polymenidou M, Rousseau F, et al. Protein Phase Separation: A New Phase in Cell Biology. Trends Cell Biol. 2018;28:420–35. 10.1016/j.tcb.2018.02.004

7. Roden C, Gladfelter AS. RNA contributions to the form and function of biomolecular condensates. Nat Rev Mol Cell Biol. 2021;22:183–95. 10.1038/s41580-020-0264-6

8. Garcia-Jove Navarro M, Kashida S, Chouaib R, Souquere S, Pierron G, Weil D, et al. RNA is a critical element for the sizing and the composition of phase-separated RNA-protein condensates. Nat Commun. 2019;10:3230. 10.1038/s41467-019-11241-6

9. Williams TD, Rousseau A. Translation regulation in response to stress. FEBS J. 2024;291:5102–22. 10.1111/febs.17076

10. Anderson P, Kedersha N. Stress granules. Curr Biol. 2009;19:R397-398. 10.1016/j.cub.2009.03.013

11. Protter DSW, Parker R. Principles and Properties of Stress Granules. Trends Cell Biol. 2016;26:668–79. 10.1016/j.tcb.2016.05.004

12. Hofmann S, Cherkasova V, Bankhead P, Bukau B, Stoecklin G. Translation suppression promotes stress granule formation and cell survival in response to cold shock. Mol Biol Cell. 2012;23:3786–800. 10.1091/mbc.E12-04-0296

13. Himanen SV, Sistonen L. New insights into transcriptional reprogramming during cellular stress. J Cell Sci. 2019;132:jcs238402. 10.1242/jcs.238402

14. Wheeler JR, Matheny T, Jain S, Abrisch R, Parker R. Distinct stages in stress granule assembly and disassembly. Nilsen TW, editor. eLife. eLife Sciences Publications, Ltd; 2016;5:e18413. 10.7554/eLife.18413

15. Jain S, Wheeler JR, Walters RW, Agrawal A, Barsic A, Parker R. ATPase-Modulated Stress Granules Contain a Diverse Proteome and Substructure. Cell. 2016;164:487–98. 10.1016/j.cell.2015.12.038

16. Sheinberger J, Shav-Tal Y. mRNPs meet stress granules. FEBS Lett. 2017;591:2534–42. 10.1002/1873-3468.12765

17. Molliex A, Temirov J, Lee J, Coughlin M, Kanagaraj AP, Kim HJ, et al. Phase Separation by Low Complexity Domains Promotes Stress Granule Assembly and Drives Pathological Fibrillization. Cell. Elsevier; 2015;163:123–33. 10.1016/j.cell.2015.09.015

18. Desai M, Gulati K, Agrawal M, Ghumra S, Sahoo PK. Stress granules: Guardians of cellular health and triggers of disease. Neural Regen Res. 2026;21:588–97. 10.4103/NRR.NRR-D-24-01196

19. Wolozin B, Ivanov P. Stress granules and neurodegeneration. Nat Rev Neurosci. 2019;20:649–66. 10.1038/s41583-019-0222-5

20. Advani VM, Ivanov P. Stress granule subtypes: an emerging link to neurodegeneration. Cell Mol Life Sci. 2020;77:4827–45. 10.1007/s00018-020-03565-0

21. Ishigaki S, Sobue G. Importance of Functional Loss of FUS in FTLD/ALS. Front Mol Biosci. 2018;5:44. 10.3389/fmolb.2018.00044

22. Guerrero EN, Wang H, Mitra J, Hegde PM, Stowell SE, Liachko NF, et al. TDP-43/FUS in motor neuron disease: Complexity and challenges. Prog Neurobiol. 2016;145–146:78–97. 10.1016/j.pneurobio.2016.09.004

23. Mariani D, Setti A, Castagnetti F, Vitiello E, Stufera Mecarelli L, Di Timoteo G, et al. ALS-associated FUS mutation reshapes the RNA and protein composition of stress granules. Nucleic Acids Res. 2024;52:13269–89. 10.1093/nar/gkae942

24. Volk AE, Weishaupt JH, Andersen PM, Ludolph AC, Kubisch C. Current knowledge and recent insights into the genetic basis of amyotrophic lateral sclerosis. Med Genet. 2018;30:252–8. 10.1007/s11825-018-0185-3

25. Sabatelli M, Moncada A, Conte A, Lattante S, Marangi G, Luigetti M, et al. Mutations in the 3′ untranslated region of FUS causing FUS overexpression are associated with amyotrophic lateral sclerosis. Hum Mol Genet. 2013;22:4748–55. 10.1093/hmg/ddt328

26. Goldstein O, Inbar T, Kedmi M, Gana-Weisz M, Abramovich B, Orr-Urtreger A, et al. FUS-P525L Juvenile Amyotrophic Lateral Sclerosis and Intellectual Disability: Evidence for Association and Oligogenic Inheritance. Neurol Genet. 2022;8:e200009. 10.1212/NXG.0000000000200009

27. Onoguchi-Mizutani R, Akimitsu N. Long noncoding RNA and phase separation in cellular stress response. J Biochem. 2022;171:269–76. 10.1093/jb/mvab156

28. Scholda J, Nguyen TTA, Kopp F. Long noncoding RNAs as versatile molecular regulators of cellular stress response and homeostasis. Hum Genet. 2024;143:813–29. 10.1007/s00439-023-02604-7

29. Aillaud M, Schulte LN. Emerging Roles of Long Noncoding RNAs in the Cytoplasmic Milieu. Noncoding RNA. 2020;6:44. 10.3390/ncrna6040044

30. Tani H. Biomolecules Interacting with Long Noncoding RNAs. Biology (Basel). 2025;14:442. 10.3390/biology14040442

31. Nishimoto Y, Nakagawa S, Okano H. NEAT1 lncRNA and amyotrophic lateral sclerosis. Neurochem Int. 2021;150:105175. 10.1016/j.neuint.2021.105175

32. Wang C, Duan Y, Duan G, Wang Q, Zhang K, Deng X, et al. Stress Induces Dynamic, Cytotoxicity-Antagonizing TDP-43 Nuclear Bodies via Paraspeckle LncRNA NEAT1-Mediated Liquid-Liquid Phase Separation. Mol Cell. 2020;79:443–458.e7. 10.1016/j.molcel.2020.06.019

33. McCluggage F, Fox AH. Paraspeckle nuclear condensates: Global sensors of cell stress? Bioessays. 2021;43:e2000245. 10.1002/bies.202000245

34. Xu Q, Liu D, Zhu L-Q, Su Y, Huang H-Z. Long non-coding RNAs as key regulators of neurodegenerative protein aggregation. Alzheimers Dement. 2025;21:e14498. 10.1002/alz.14498

35. Vangoor VR, Gomes-Duarte A, Pasterkamp RJ. Long non-coding RNAs in motor neuron development and disease. J Neurochem. 2021;156:777–801. 10.1111/jnc.15198

36. Falduti A, Giovinazzo A, Lo Feudo E, Rocca V, Brighina F, Messina A, et al. The Role of Non-Coding RNAs in ALS. Genes. Multidisciplinary Digital Publishing Institute; 2025;16:623. 10.3390/genes16060623

37. Dini Modigliani S, Morlando M, Errichelli L, Sabatelli M, Bozzoni I. An ALS-associated mutation in the FUS 3′-UTR disrupts a microRNA–FUS regulatory circuitry. Nat Commun. Nature Publishing Group; 2014;5:4335. 10.1038/ncomms5335

38. Garone MG, de Turris V, Soloperto A, Brighi C, De Santis R, Pagani F, et al. Conversion of Human Induced Pluripotent Stem Cells (iPSCs) into Functional Spinal and Cranial Motor Neurons Using PiggyBac Vectors. J Vis Exp. 2019; 10.3791/59321

39. Choi HMT, Schwarzkopf M, Fornace ME, Acharya A, Artavanis G, Stegmaier J, et al. Third-generation in situ hybridization chain reaction: multiplexed, quantitative, sensitive, versatile, robust. Development. Cambridge, England; 2018;145:dev165753. 10.1242/dev.165753

40. Buonaiuto G, Santini T, Ballarino M. Protocol for the simultaneous detection of nuclear long non-coding RNAs and proteins in human iPSC-derived cardiomyocytes. STAR Protoc. 2026;7:104351. 10.1016/j.xpro.2026.104351

41. Zhang Y, Pak C, Han Y, Ahlenius H, Zhang Z, Chanda S, et al. Rapid Single-Step Induction of Functional Neurons from Human Pluripotent Stem Cells. Neuron. 2013;78:785–98. 10.1016/j.neuron.2013.05.029

42. Martin M. Cutadapt removes adapter sequences from high-throughput sequencing reads. EMBnet.journal. 2011;17:10–2. 10.14806/ej.17.1.200

43. Bolger AM, Lohse M, Usadel B. Trimmomatic: a flexible trimmer for Illumina sequence data. Bioinformatics. 2014;30:2114–20. 10.1093/bioinformatics/btu170

44. Langmead B, Salzberg SL. Fast gapped-read alignment with Bowtie 2. Nat Methods. Nature Publishing Group; 2012;9:357–9. 10.1038/nmeth.1923

45. Dobin A, Davis CA, Schlesinger F, Drenkow J, Zaleski C, Jha S, et al. STAR: ultrafast universal RNA-seq aligner. Bioinformatics. 2013;29:15–21. 10.1093/bioinformatics/bts635

46. Li H, Handsaker B, Wysoker A, Fennell T, Ruan J, Homer N, et al. The Sequence Alignment/Map format and SAMtools. Bioinformatics. 2009;25:2078–9. 10.1093/bioinformatics/btp352

47. Anders S, Pyl PT, Huber W. HTSeq—a Python framework to work with high-throughput sequencing data. Bioinformatics. 2015;31:166–9. 10.1093/bioinformatics/btu638

48. Yates AD, Achuthan P, Akanni W, Allen J, Allen J, Alvarez-Jarreta J, et al. Ensembl 2020. Nucleic Acids Res. 2020;48:D682–8. 10.1093/nar/gkz966

49. Robinson MD, McCarthy DJ, Smyth GK. edgeR: a Bioconductor package for differential expression analysis of digital gene expression data. Bioinformatics. 2010;26:139–40. 10.1093/bioinformatics/btp616

50. De Santis R, Santini L, Colantoni A, Peruzzi G, de Turris V, Alfano V, et al. FUS Mutant Human Motoneurons Display Altered Transcriptome and microRNA Pathways with Implications for ALS Pathogenesis. Stem Cell Reports. 2017;9:1450–62. 10.1016/j.stemcr.2017.09.004

51. Patro R, Duggal G, Love MI, Irizarry RA, Kingsford C. Salmon provides fast and bias-aware quantification of transcript expression. Nat Methods. Nature Publishing Group; 2017;14:417–9. 10.1038/nmeth.4197

52. Basu S, Rajendra KC, Alagar S, Bahadur RP. Impaired nuclear transport induced by juvenile ALS causing P525L mutation in NLS domain of FUS: A molecular mechanistic study. Biochim Biophys Acta Proteins Proteom. 2022;1870:140766. 10.1016/j.bbapap.2022.140766

53. Lo Bello M, Di Fini F, Notaro A, Spataro R, Conforti FL, La Bella V. ALS-Related Mutant FUS Protein Is Mislocalized to Cytoplasm and Is Recruited into Stress Granules of Fibroblasts from Asymptomatic FUS P525L Mutation Carriers. Neurodegener Dis. 2017;17:292–303. 10.1159/000480085

54. Murakami T, Qamar S, Lin JQ, Schierle GSK, Rees E, Miyashita A, et al. ALS/FTD Mutation-Induced Phase Transition of FUS Liquid Droplets and Reversible Hydrogels into Irreversible Hydrogels Impairs RNP Granule Function. Neuron. 2015;88:678–90. 10.1016/j.neuron.2015.10.030

55. Di Timoteo G, Giuliani A, Setti A, Biagi MC, Lisi M, Santini T, et al. M6A reduction relieves FUS-associated ALS granules. Nat Commun. 2024;15:5033. 10.1038/s41467-024-49416-5

56. Khong A, Matheny T, Jain S, Mitchell SF, Wheeler JR, Parker R. The Stress Granule Transcriptome Reveals Principles of mRNA Accumulation in Stress Granules. Mol Cell. 2017;68:808–820.e5. 10.1016/j.molcel.2017.10.015

57. Lenzi J, De Santis R, de Turris V, Morlando M, Laneve P, Calvo A, et al. ALS mutant FUS proteins are recruited into stress granules in induced pluripotent stem cell-derived motoneurons. Dis Model Mech. 2015;8:755–66. 10.1242/dmm.020099

58. Yang P, Mathieu C, Kolaitis R-M, Zhang P, Messing J, Yurtsever U, et al. G3BP1 Is a Tunable Switch that Triggers Phase Separation to Assemble Stress Granules. Cell. 2020;181:325–345.e28. 10.1016/j.cell.2020.03.046

59. Kim JB, Greber B, Araúzo-Bravo MJ, Meyer J, Park KI, Zaehres H, et al. Direct reprogramming of human neural stem cells by OCT4. Nature. Nature Publishing Group; 2009;461:649–53. 10.1038/nature08436

60. Arber S, Han B, Mendelsohn M, Smith M, Jessell TM, Sockanathan S. Requirement for the homeobox gene Hb9 in the consolidation of motor neuron identity. Neuron. 1999;23:659–74. 10.1016/s0896-6273(01)80026-x

61. Liang X, Song M-R, Xu Z, Lanuza GM, Liu Y, Zhuang T, et al. Isl1 is required for multiple aspects of motor neuron development. Mol Cell Neurosci. 2011;47:215–22. 10.1016/j.mcn.2011.04.007

62. Patel A, Lee HO, Jawerth L, Maharana S, Jahnel M, Hein MY, et al. A Liquid-to-Solid Phase Transition of the ALS Protein FUS Accelerated by Disease Mutation. Cell. 2015;162:1066–77. 10.1016/j.cell.2015.07.047

63. Chujo T, Hirose T. Nuclear Bodies Built on Architectural Long Noncoding RNAs: Unifying Principles of Their Construction and Function. Mol Cells. 2017;40:889–96. 10.14348/molcells.2017.0263

64. Yamazaki T, Nakagawa S, Hirose T. Architectural RNAs for Membraneless Nuclear Body Formation. Cold Spring Harb Symp Quant Biol. 2019;84:227–37. 10.1101/sqb.2019.84.039404

65. Fujiwara N, Ueno T, Yamazaki T, Hirose T. Unraveling architectural RNAs: Structural and functional blueprints of membraneless organelles and strategies for genome-scale identification. Biochim Biophys Acta Gen Subj. 2025;1869:130815. 10.1016/j.bbagen.2025.130815

66. Gellatly SA, Kalujnaia S, Cramb G. Cloning, tissue distribution and sub-cellular localisation of phospholipase C X-domain containing protein (PLCXD) isoforms. Biochem Biophys Res Commun. 2012;424:651–6. 10.1016/j.bbrc.2012.06.079

67. Morin GM, Zerbib L, Kaltenbach S, Fraissenon A, Balducci E, Asnafi V, et al. PIK3CA-Related Disorders: From Disease Mechanism to Evidence-Based Treatments. Annu Rev Genomics Hum Genet. 2024;25:211–37. 10.1146/annurev-genom-121222-114518

68. Zhen Y, Stenmark H. Cellular functions of Rab GTPases at a glance. J Cell Sci. 2015;128:3171–6. 10.1242/jcs.166074

69. Aljaibeji H, Mukhopadhyay D, Mohammed AK, Dhaiban S, Hachim MY, Elemam NM, et al. Reduced Expression of PLCXD3 Associates With Disruption of Glucose Sensing and Insulin Signaling in Pancreatic β-Cells. Front Endocrinol (Lausanne). 2019;10:735. 10.3389/fendo.2019.00735

70. Koundouros N, Poulogiannis G. Phosphoinositide 3-Kinase/Akt Signaling and Redox Metabolism in Cancer. Front Oncol. 2018;8:160. 10.3389/fonc.2018.00160

71. Zhang Y, Zhao J, Chen X, Qiao Y, Kang J, Guo X, et al. DHX36 binding induces RNA structurome remodeling and regulates RNA abundance via m6A reader YTHDF1. Nat Commun. Nature Publishing Group; 2024;15:9890. 10.1038/s41467-024-54000-y

72. Antcliff A, McCullough LD, Tsvetkov AS. G-Quadruplexes and the DNA/RNA helicase DHX36 in health, disease, and aging. Aging (Albany NY). 2021;13:25578–87. 10.18632/aging.203738

73. Lu L, Zheng L, Si Y, Luo W, Dujardin G, Kwan T, et al. Hu antigen R (HuR) is a positive regulator of the RNA-binding proteins TDP-43 and FUS/TLS: implications for amyotrophic lateral sclerosis. J Biol Chem. 2014;289:31792–804. 10.1074/jbc.M114.573246

74. Good PJ. A conserved family of elav-like genes in vertebrates. Proc Natl Acad Sci U S A. 1995;92:4557–61. 10.1073/pnas.92.10.4557

75. Gallouzi IE, Brennan CM, Stenberg MG, Swanson MS, Eversole A, Maizels N, et al. HuR binding to cytoplasmic mRNA is perturbed by heat shock. Proc Natl Acad Sci U S A. 2000;97:3073–8. 10.1073/pnas.97.7.3073

76. Kedersha NL, Gupta M, Li W, Miller I, Anderson P. RNA-Binding Proteins Tia-1 and Tiar Link the Phosphorylation of Eif-2α to the Assembly of Mammalian Stress Granules. J Cell Biol. 1999;147:1431–42. 10.1083/jcb.147.7.1431

77. Waris S, Wilce MCJ, Wilce JA. RNA recognition and stress granule formation by TIA proteins. Int J Mol Sci. 2014;15:23377–88. 10.3390/ijms151223377

78. Jeon P, Ham H-J, Park S, Lee J-A. Regulation of Cellular Ribonucleoprotein Granules: From Assembly to Degradation via Post-translational Modification. Cells. Multidisciplinary Digital Publishing Institute; 2022;11:2063. 10.3390/cells11132063

79. Liao J-Y, Yang B, Shi C-P, Deng W-X, Deng J-S, Cen M-F, et al. RBPWorld for exploring functions and disease associations of RNA-binding proteins across species. Nucleic Acids Res. 2025;53:D220–32. 10.1093/nar/gkae1028

80. Schult P, Paeschke K. The DEAH helicase DHX36 and its role in G-quadruplex-dependent processes. Biol Chem. 2021;402:581–91. 10.1515/hsz-2020-0292

81. Kuwano Y, Gorospe M. Protecting the stress response, guarding the MKP-1 mRNA. Cell Cycle. Taylor & Francis; 2008;7:2640–2. 10.4161/cc.7.17.6534

82. Gorospe M. HuR in the Mammalian Genotoxic Response: Post-Transcriptional Multitasking. Cell Cycle. Taylor & Francis; 2003;2:411–3. 10.4161/cc.2.5.491

83. Nishimoto Y, Nakagawa S, Hirose T, Okano HJ, Takao M, Shibata S, et al. The long non-coding RNA nuclear-enriched abundant transcript 1_2 induces paraspeckle formation in the motor neuron during the early phase of amyotrophic lateral sclerosis. Mol Brain. 2013;6:31. 10.1186/1756-6606-6-31

84. Elguindy MM, Mendell JT. NORAD-induced PUMILIO phase separation is required for genome stability. Nature. 2021;595:303–8. 10.1038/s41586-021-03633-w

85. Lee S, Kopp F, Chang T-C, Sataluri A, Chen B, Sivakumar S, et al. Noncoding RNA NORAD Regulates Genomic Stability by Sequestering PUMILIO Proteins. Cell. 2016;164:69–80. 10.1016/j.cell.2015.12.017

