## Supplementary figure legends for "*LINC00205* acts as a multivalent scaffold promoting FUS^P525L^ recruitment in Amyotrophic Lateral Sclerosis stress granules"

**Supplementary Fig.1: *LINC00205* is enriched in SG and interacts with FUS$^{P525L}$. A)** *LINC00205* splice variants from Ensembl database. **B)** Histogram showing expression of *LINC00205* isoforms in MN (DAY 12), MN (DAY 12+7) and SK-N-BE (DOX-untreated, ARS-treated). Data are shown as transcript per million (TPM) $\pm$ SEM. N=3 for MN (DAY 12 and DAY 12+7); N=4 for SK-N-BE. **C)** qRT-PCR analysis of *LINC00205* enrichment over INP in SG purified from ARS-treated FUS$^{P525L}$ SK-N-BE cells. SG from untreated cells served as negative control. Data are expressed as mean INP percentage ± SEM. *HUWE1* and *ATP5O* served as positive and negative controls, respectively. N=3. **D)** qRT-PCR analysis of *LINC00205* enrichment over INP in SG purified from ARS-treated FUS$^{WT}$ SK-N-BE cells. Details as in C. **E)** Representative western blot of FUS$^{P525L}$ levels in INP 10% and FLAG-FUS$^{P525L}$ CLIP samples from untreated FUS$^{P525L}$ SK-N-BE cells, DOX-treated (+DOX). DOX-untreated (-DOX) cells served as negative control. GAPDH served as loading control. **F)** Line graphs showing *LINC00205* expression during neuronal differentiation from FUS$^{WT}$ (left) and FUS$^{P525L}$ (right) hiPSC to MN (DAY0, DAY5, DIV3 and DIV7). Data are expressed as mean ± SEM (arbitrary units, A.U.) of relative quantities normalized to *ATP5O*. N=2. **G)** Representative smFISH/immunofluorescence confocal images of ARS-treated FUS$^{P525L}$ MN showing colocalization of *LINC00205* (left panel) or *GAPDH* (right panel) with G3BP1-positive SG. FUS staining marks nuclei. **H)** Boxplot quantifying *LINC00205*/G3BP1-positive SG colocalization in ARS-treated FUS$^{WT}$ MN. *GAPDH* served as negative control. 222 and 193 cells were analyzed for *LINC00205* and *GAPDH* colocalization, respectively. N=2. ****$p \leq 0.0001$ (two-tailed, unpaired Student's T-test). **I)** Representative smFISH/immunofluorescence confocal images of ARS-treated

FUS$^{WT}$ MN showing colocalization of *LINC00205* (left) or *GAPDH* (right) with G3BP1-positive SG. FUS staining marks nuclei. **J)** Representative smFISH/immunofluorescence confocal images of ARS-treated FUS$^{WT}$ MN showing colocalization of *LINC00205* (upper-left) and *GAPDH* (lower-left) with G3BP1-positive SG. FUS staining marks nuclei. 3D renderings (upper and lower-right) are derived from indicated regions.

**Supplementary Fig.2: *LINC00205* depletion reduces the number of FUS$^{P525L}$-containing SG in human MN. A)** CRISPR/Cas9 experiment design in FUS$^{WT}$ and FUS$^{P525L}$ hiPSC. A cassette containing a Poly-adenylation sequence (PAS) and Neomycin resistance (NeoR) was inserted downstream of *LINC00205* exon 1 via Cas9 and two sgRNAs. Two homology arms complementary to sequences upstream and downstream of the cut site were used. **B)** qRT-PCR analysis of *LINC00205* expression in *LINC00205* KOs FUS$^{WT}$ hiPSC (wt-KO#1 and wt-KO#2, left) and FUS$^{P525L}$ (525-KO#1 and 525-KO#2, right) compared to controls, set as 1. Data are expressed as mean ± SEM (arbitrary units, A.U.) of fold changes normalized to *ATP5O*. N=3. ****$p \leq 0.0001$ (two-tailed, paired Student's T-test). **C-D)** Line graphs showing expression of differentiation markers *OCT4* (left panel) *HB9* (middle panel)*, ISL1* (right panel) during differentiation from FUS$^{WT}$ (C) and FUS$^{P525L}$ (D) hiPSC to MN (DAY0, DAY5, DIV3 and DIV7). Data are expressed as mean ± SEM (arbitrary units, A.U.) of relative quantities normalized to *ATP5O*. N=2. n.s. $p > 0.05$ (two-tailed, paired Student's T-test). Comparisons were performed between CTRL and KO#1 or KO#2 at each time point. **E)** Representative immunofluorescence confocal images of ARS-treated FUS$^{WT}$ MN showing G3BP1-positive SG in wt-CTRL (upper row), wt-KO#1 (middle row) and wt-KO#2 (lower row) cells. Nuclei were counterstained with DAPI. Right column panels show magnifications of the indicated regions. **F-I)** FUS expression analysis in *LINC00205* KO compared to CTRL $FUS^{P525L}$ (F-G) and $FUS^{WT}$ (H-I) MN untreated or ARS-treated. Left: qRT-PCR analysis of *FUS* expression levels. Data are expressed as mean ± SEM (arbitrary units, A.U.) of relative quantities normalized to *ATP5O*. N=3. Right: representative western blot of FUS protein levels. In the histogram below, FUS protein levels were quantified relative to GAPDH, setting the CTRL as 1. Data are expressed as means ± SEM (arbitrary units, A.U.). N=3. n.s. p>0.05 (two-tailed, paired Student's T-test).

**Supplementary Fig.3: *LINC00205* loss favors faster SG recovery in $FUS^{P525L}$ MN. A)** Representative immunofluorescence confocal images of $FUS^{WT}$ MN at different time points (NT, 1h ARS, 2h REC, 4h REC, each column) in wt-CTRL (upper row), wt-KO#1 (middle row) and wt-KO#2 (lower row) cells. Nuclei were counterstained with DAPI.

**Supplementary Fig.4: *LINC00205* promotes selective mRNA localization to $FUS^{P525L}$-containing SG. A-B)** qRT-PCR analysis of *LINC00205* enrichment over INP in *LINC00205* and *LACZ* native RNA PD for RNA-seq performed in untreated (left panels) and ARS-treated (right panels) of $FUS^{P525L}$ (A) and $FUS^{WT}$ (B) SK-N-BE cells. Data are expressed as mean of INP percentage ± SEM. *ATP5O* served as negative control. N=2. **C)** Venn diagram summarizing the number of *LINC00205*-interacting transcripts identified from *LINC00205* native PD followed by RNA-Seq performed in untreated and ARS-treated $FUS^{WT}$ and $FUS^{P525L}$ SK-N-BE. **D)** Venn diagram showing the number of transcripts bound to *LINC00205* in ARS-treated FUS$^{WT}$ and FUS$^{P525L}$ SK-N-BE. **E-H)** qRT-PCR validation of RNA-seq results. Analysis of *LINC00205* (left panels) and *PLCXD3*, *ZNF841*, *RAB30* and *PIK3CA* mRNA (right panels) enrichment over INP in *LINC00205* and *LACZ* native RNA PD performed in FUS$^{P525L}$ SK-N-BE cells untreated (E) or ARS-treated (F), and in FUS$^{WT}$ SK-N-BE untreated (G) or ARS-treated (H). Data are expressed as mean of INP percentage ± SEM. *ATP5O* served as negative control. N=2. **I)** Representative smFISH/immunofluorescence confocal images of ARS-treated FUS$^{WT}$ and FUS$^{P525L}$ MN showing colocalization *GAPDH* with G3BP1-positive SG in wt-CTRL (upper row) 525-CTRL (middle-upper row), 525-KO#1 (middle-lower row) and 525-KO#2 (lower row) cells. FUS staining marks nuclei. 3D renderings (middle-right and right columns) are derived from indicated regions.

**Supplementary Fig.5: LINC00205 directly interacts with DHX36 and increases its colocalization with FUS$^{P525L}$-containing SG. A)** Representative western blot of DHX36 levels in INP 10%, DHX36 CLIP and IgG CLIP samples obtained from untreated FUS$^{P525L}$ SK-N-BE cells. GAPDH served as loading control. **B)** Left panel: representative western blot of HuR levels in INP 10%, HuR CLIP and IgG CLIP samples obtained from untreated FUS$^{P525L}$ SK-N-BE cells. ACTININ served as loading control. Right panel: qRT-PCR analysis of *LINC00205* enrichment over INP in IP and IgG fractions from HuR CLIP assay performed in untreated FUS$^{P525L}$ SK-N-BE cells. Data are expressed as mean of INP percentage ± SEM. *HuR* and *GAPDH* were used as positive and negative controls, respectively. N=3. **C)** Representative immunofluorescence confocal images of ARS-treated FUS$^{WT}$ MN showing DHX36/TIAR-positive SG colocalization in wt-CTRL (upper row), wt-KO#1 (middle row) and wt-KO#2 (lower row) cells. Nuclei were counterstained with DAPI. Right panels show magnifications of the indicated regions. All images represent a single focal plane from the acquired z-stacks. **D)** Boxplot showing Pearson's correlation coefficients indicating HuR/G3BP1-positive SG colocalization by immunofluorescence analyses in ARS-treated $FUS^{P525L}$ and $FUS^{WT}$ MN. For $FUS^{P525L}$ and $FUS^{WT}$ MN conditions (CTRL, KO#1, KO#2), the number of cells analyzed were 652, 320, 510 and 398, 204, 331, respectively. N=2. n.s., $p>0.05$ (two-tailed, unpaired Student's T-test). **E-F)** Representative immunofluorescence confocal images of ARS-treated $FUS^{P525L}$ (E) and $FUS^{WT}$ (F) MN showing HuR/G3BP1-positive SG colocalization in CTRL (upper row), KO#1 (middle row) and KO#2 (lower row) cells. Nuclei were counterstained with DAPI. Right panels show magnifications of the indicated regions. Images represent a single focal plane from the acquired z-stacks.
