## Supplementary figures and images for "*LINC00205* acts as a multivalent scaffold promoting FUS^P525L^ recruitment in Amyotrophic Lateral Sclerosis stress granules"

### Supplementary Fig. 1 A-E

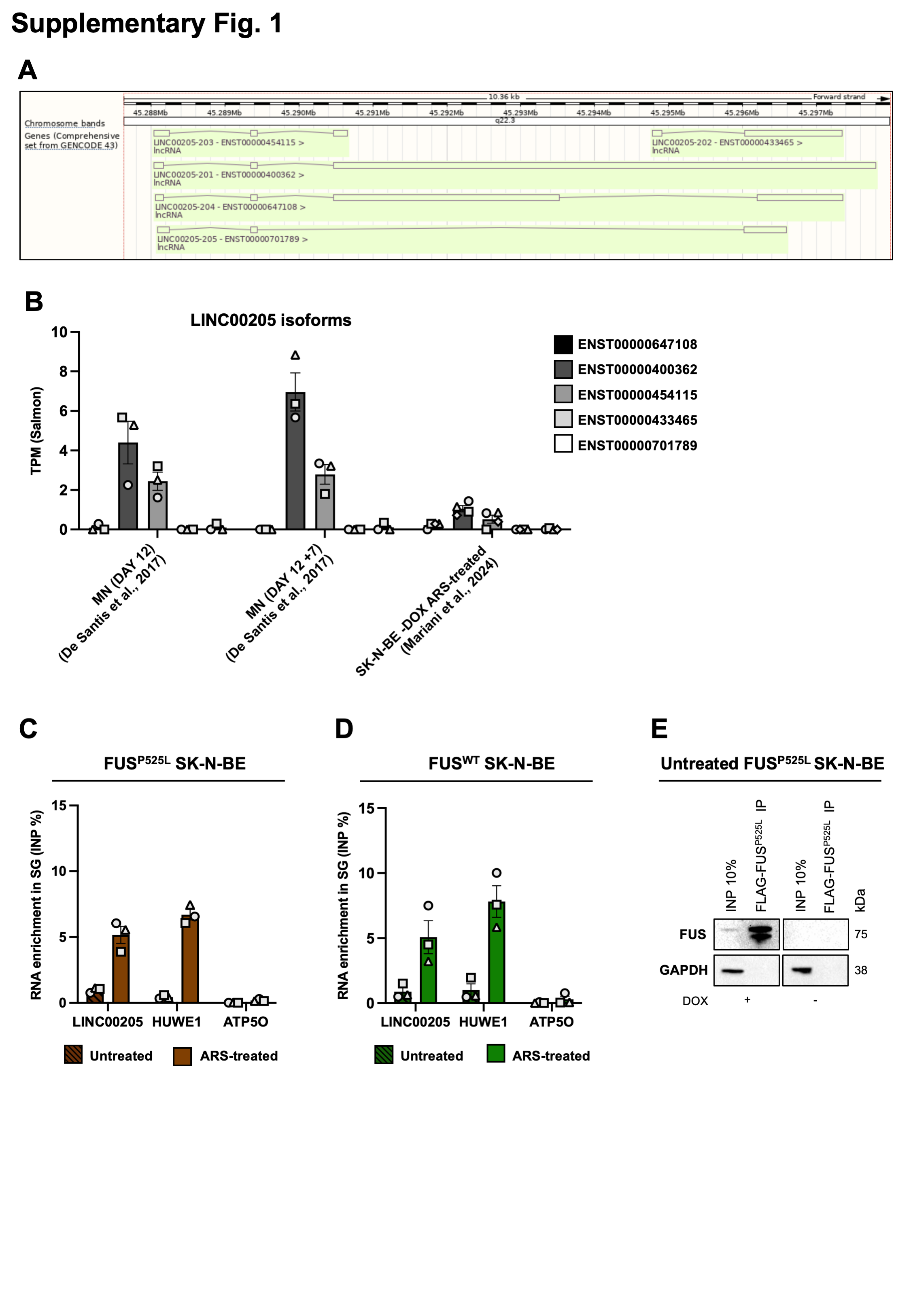

### Supplementary Fig. 1 F-G

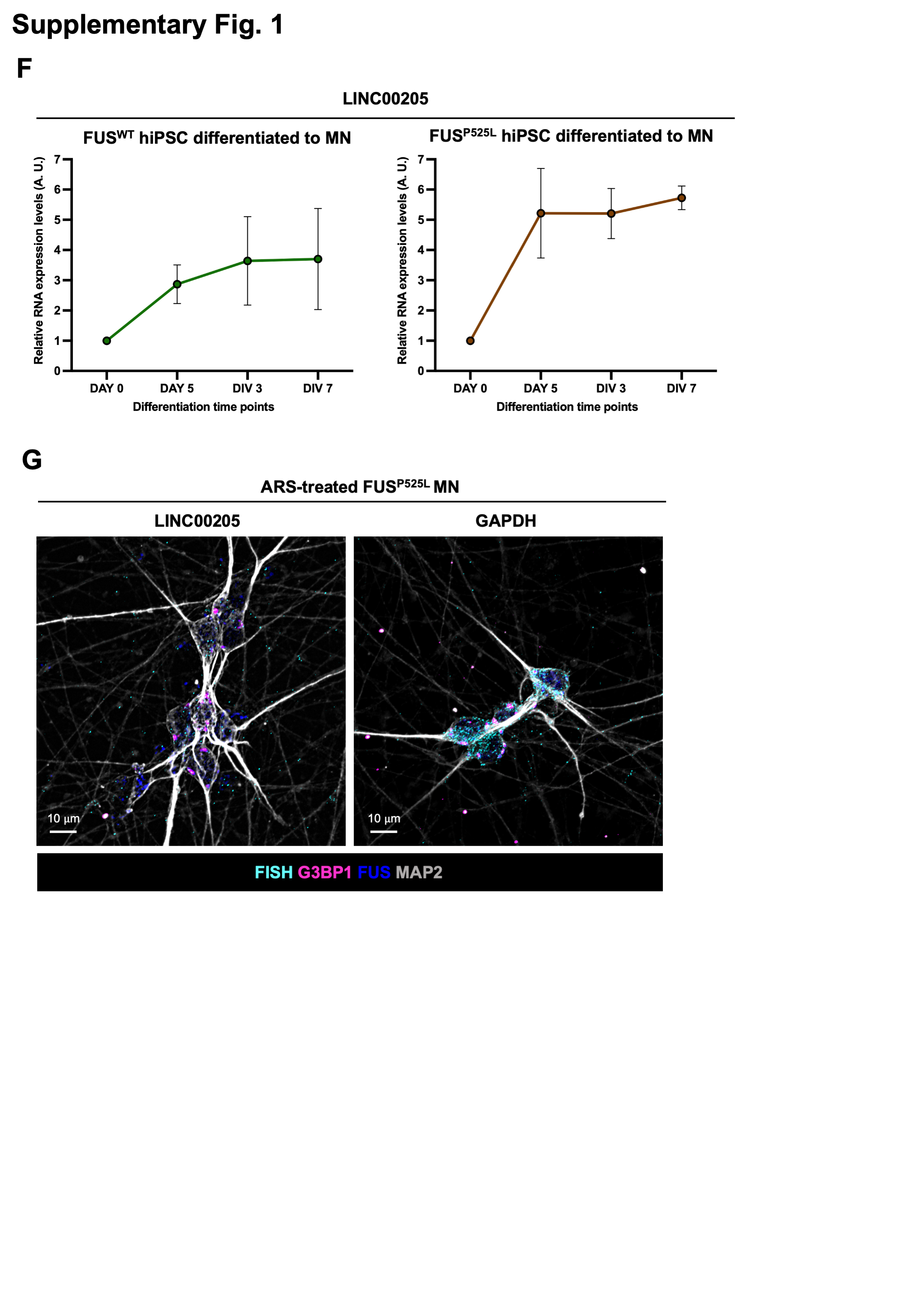

### Supplementary Fig. 1 H-J

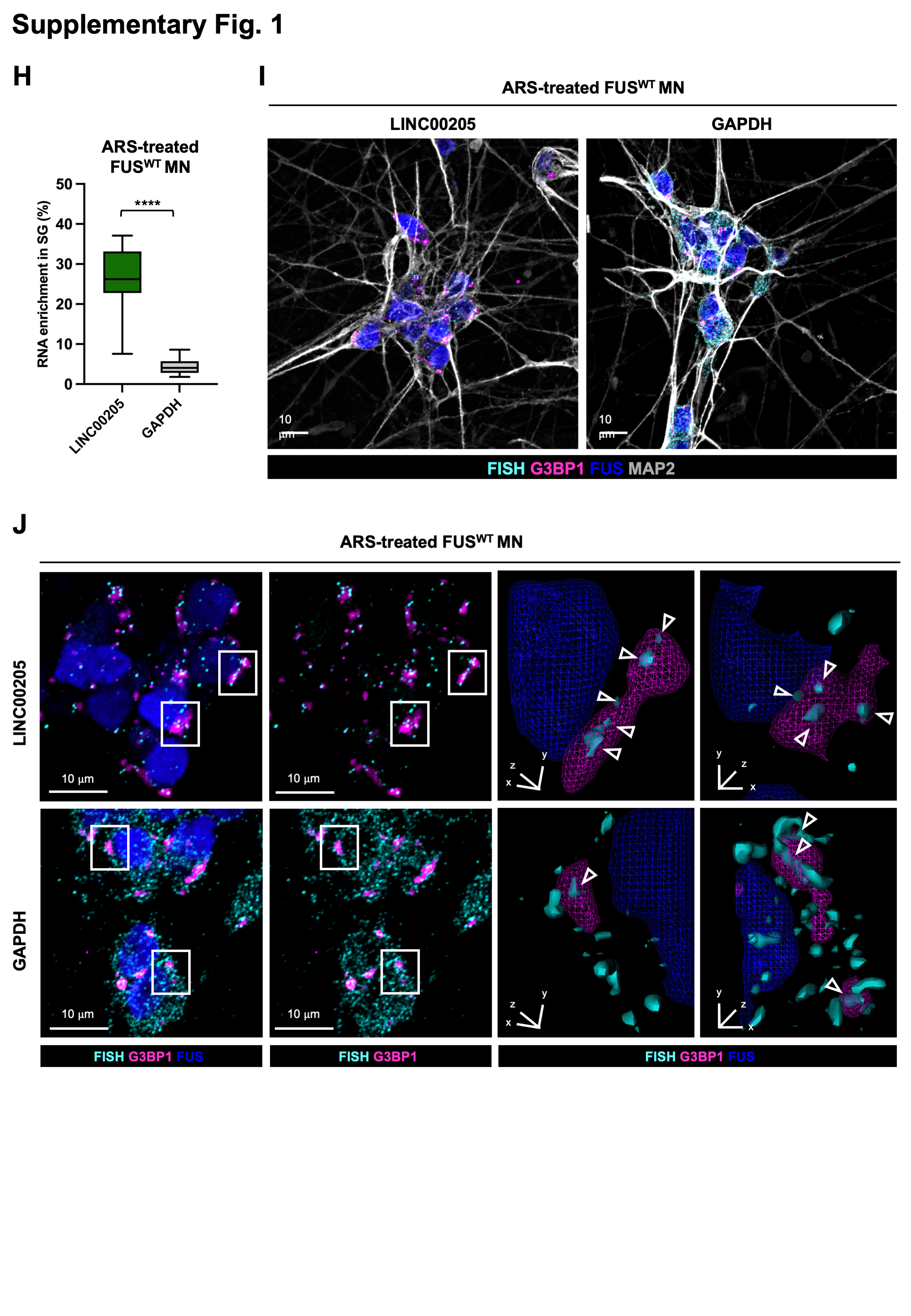

### Supplementary Fig. 2 A-D

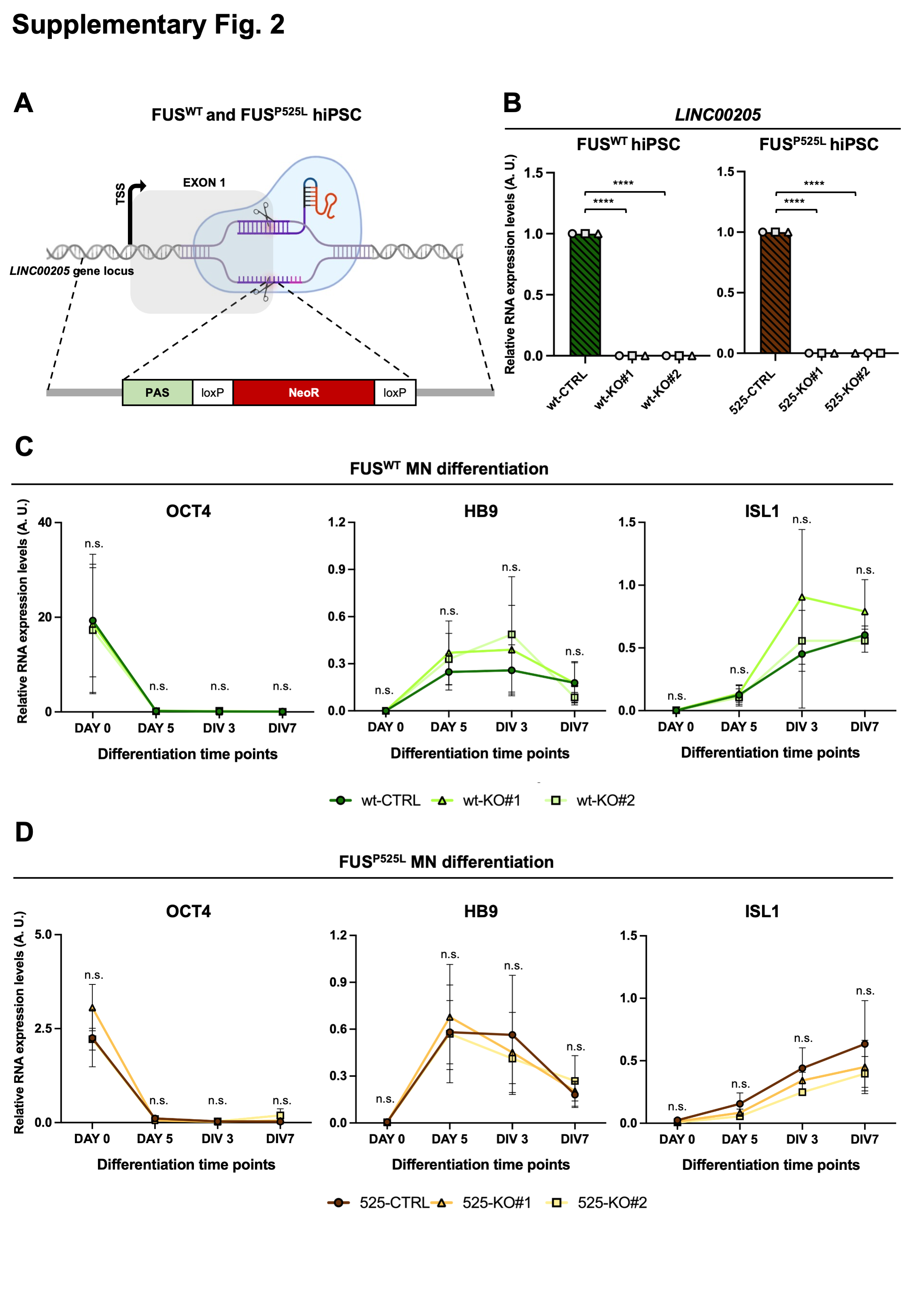

### Supplementary Fig. 2 E-I

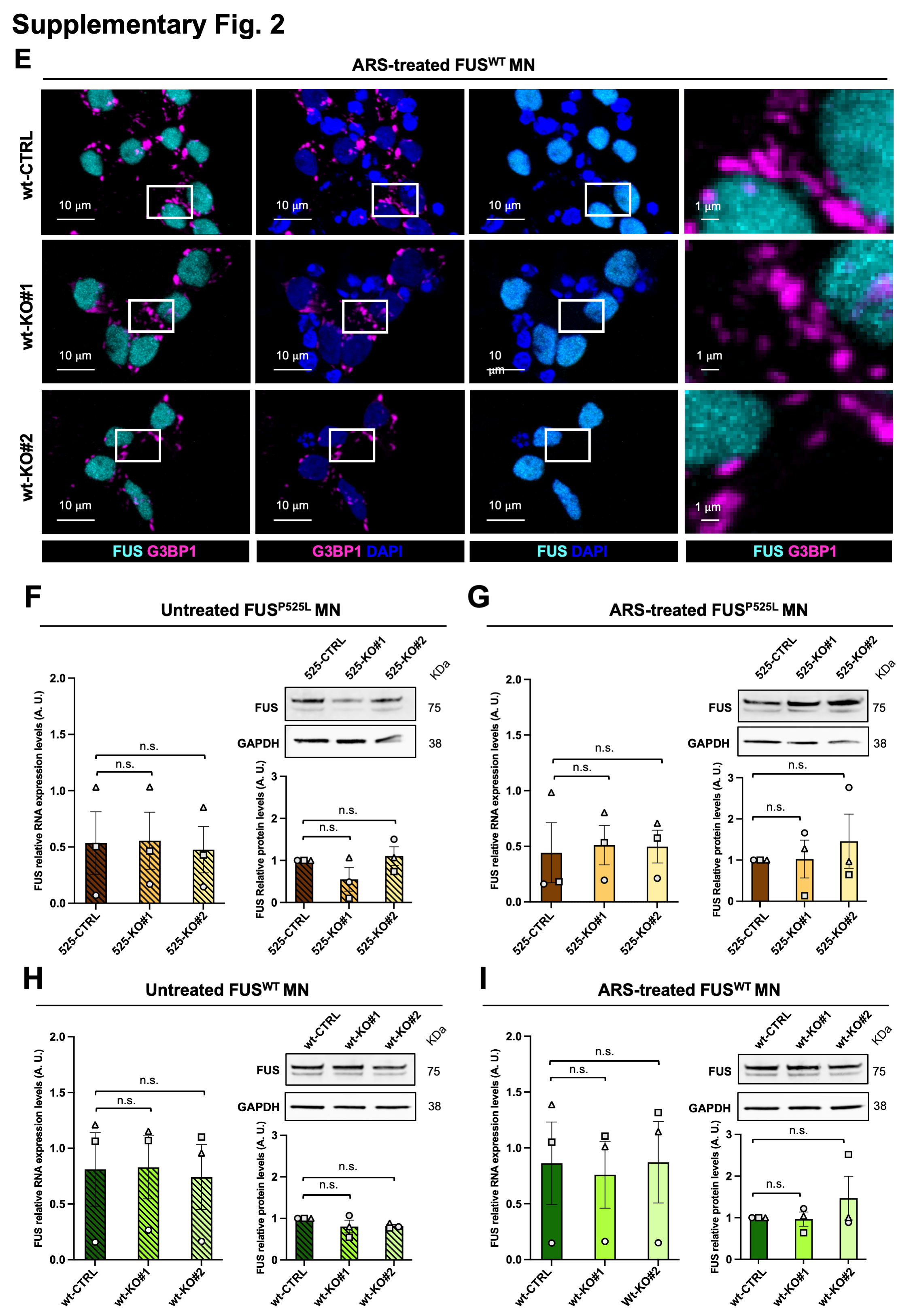

### Supplementary Fig. 3 A

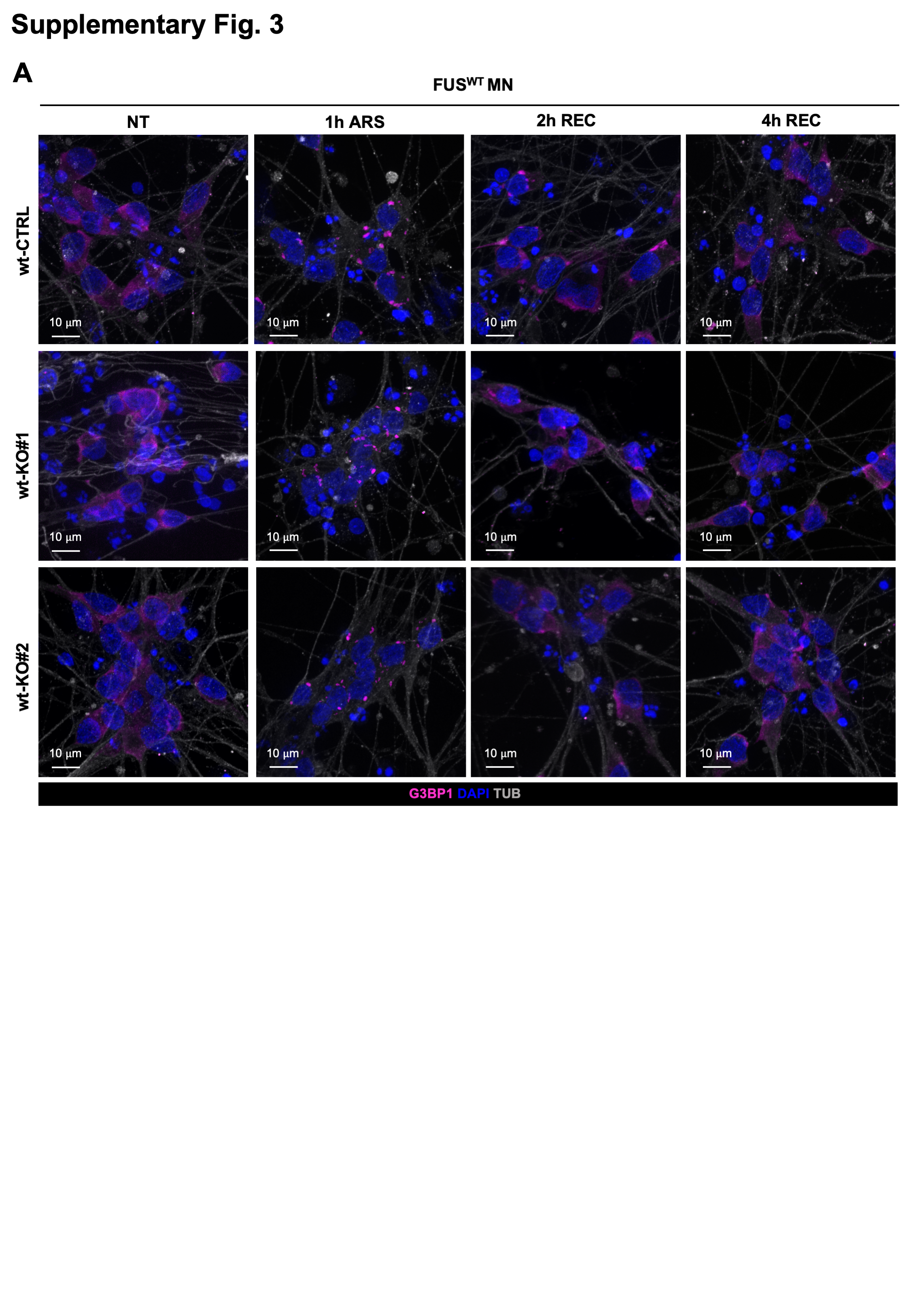

### Supplementary Fig. 4 A-H

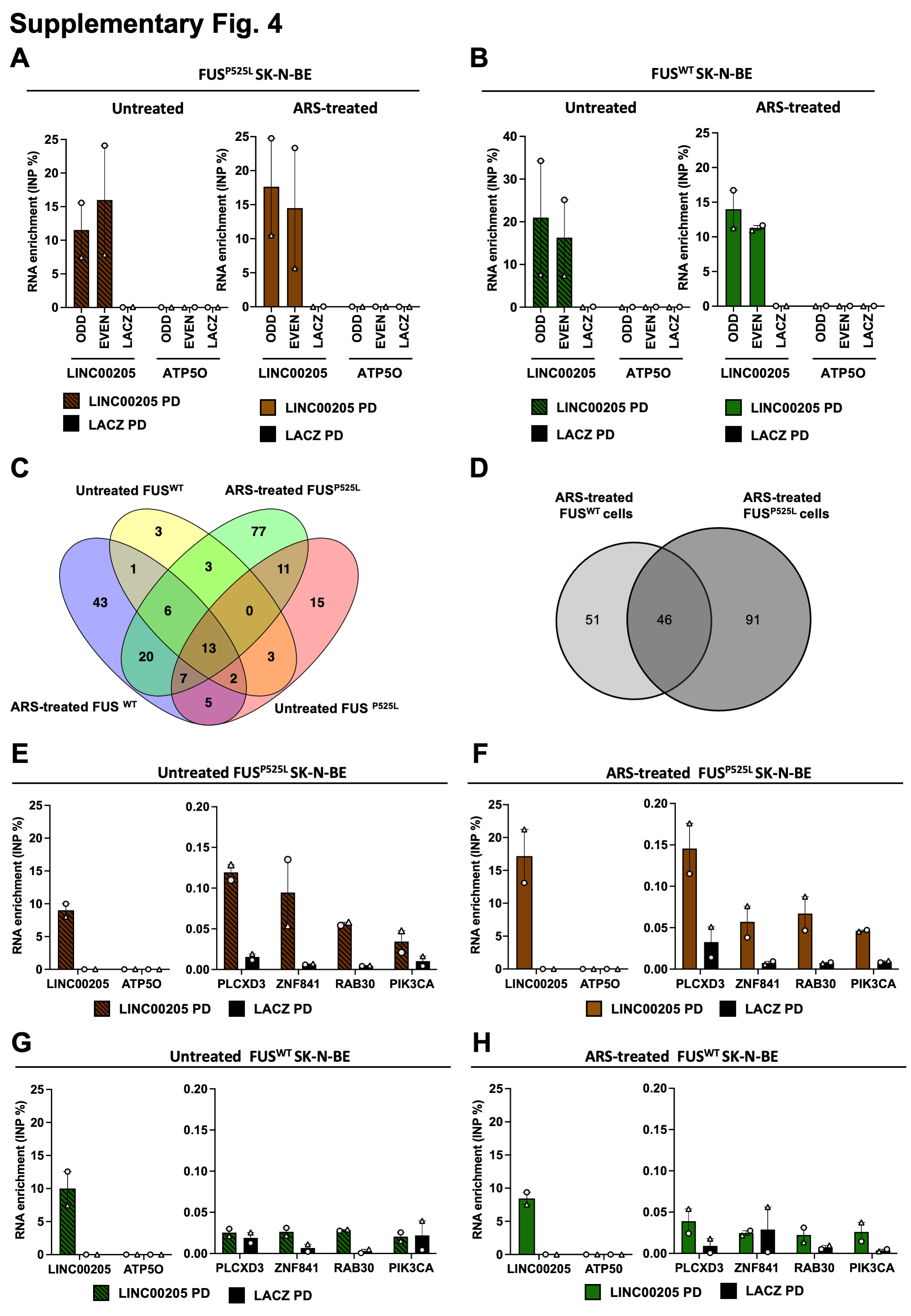

### Supplementary Fig. 4 I

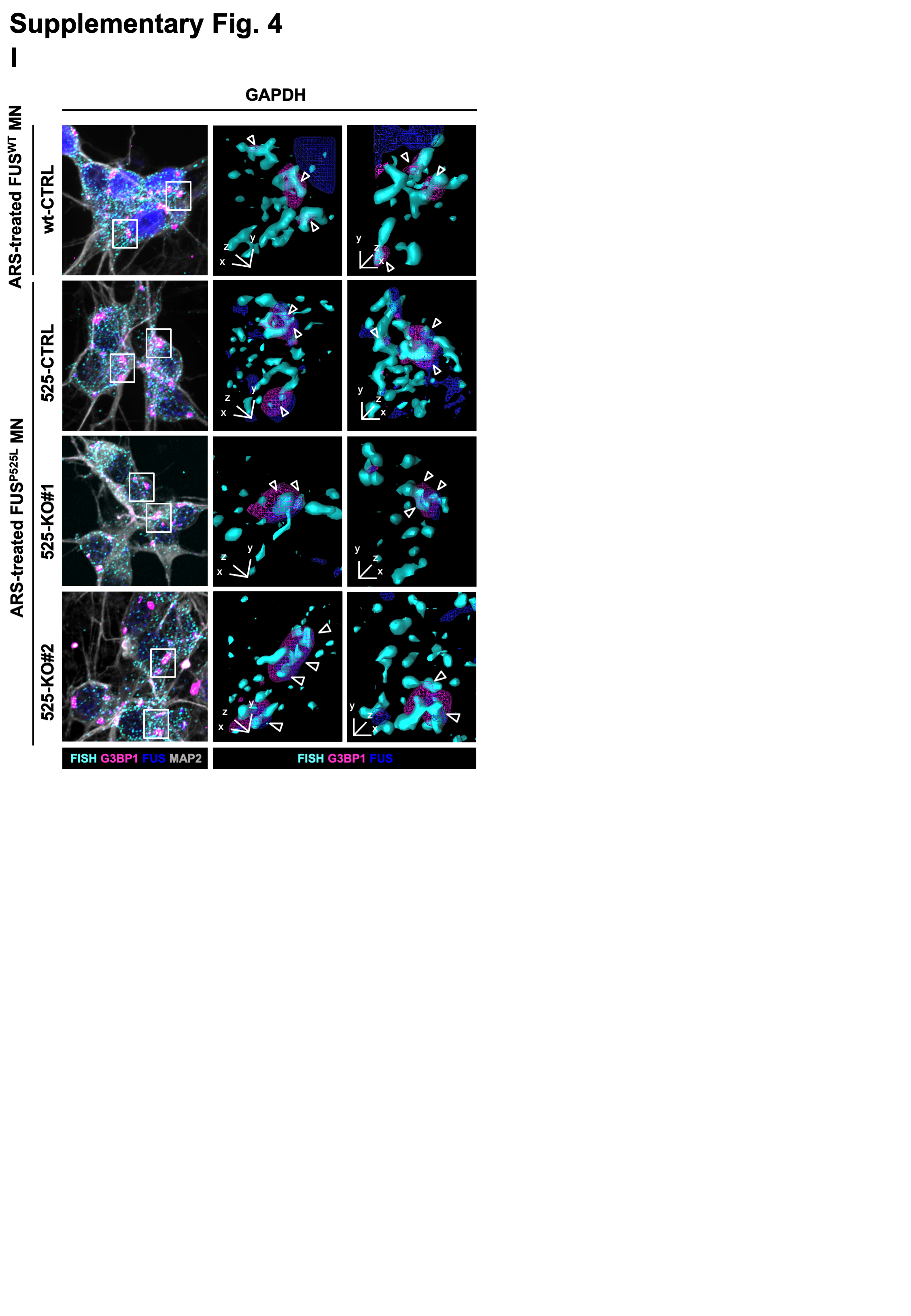

### Supplementary Fig. 5 A-C

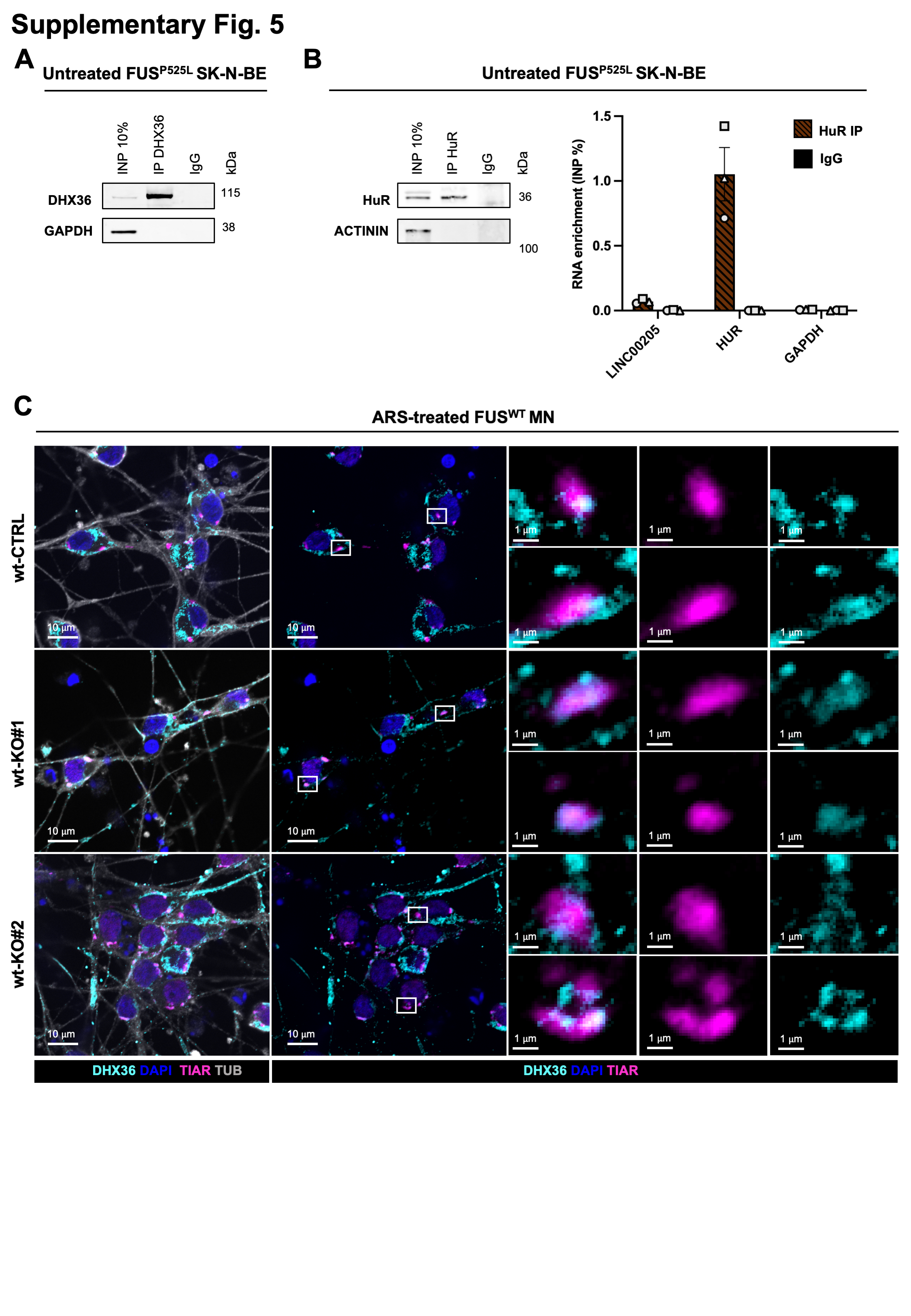

### Supplementary Fig. 5 D-E

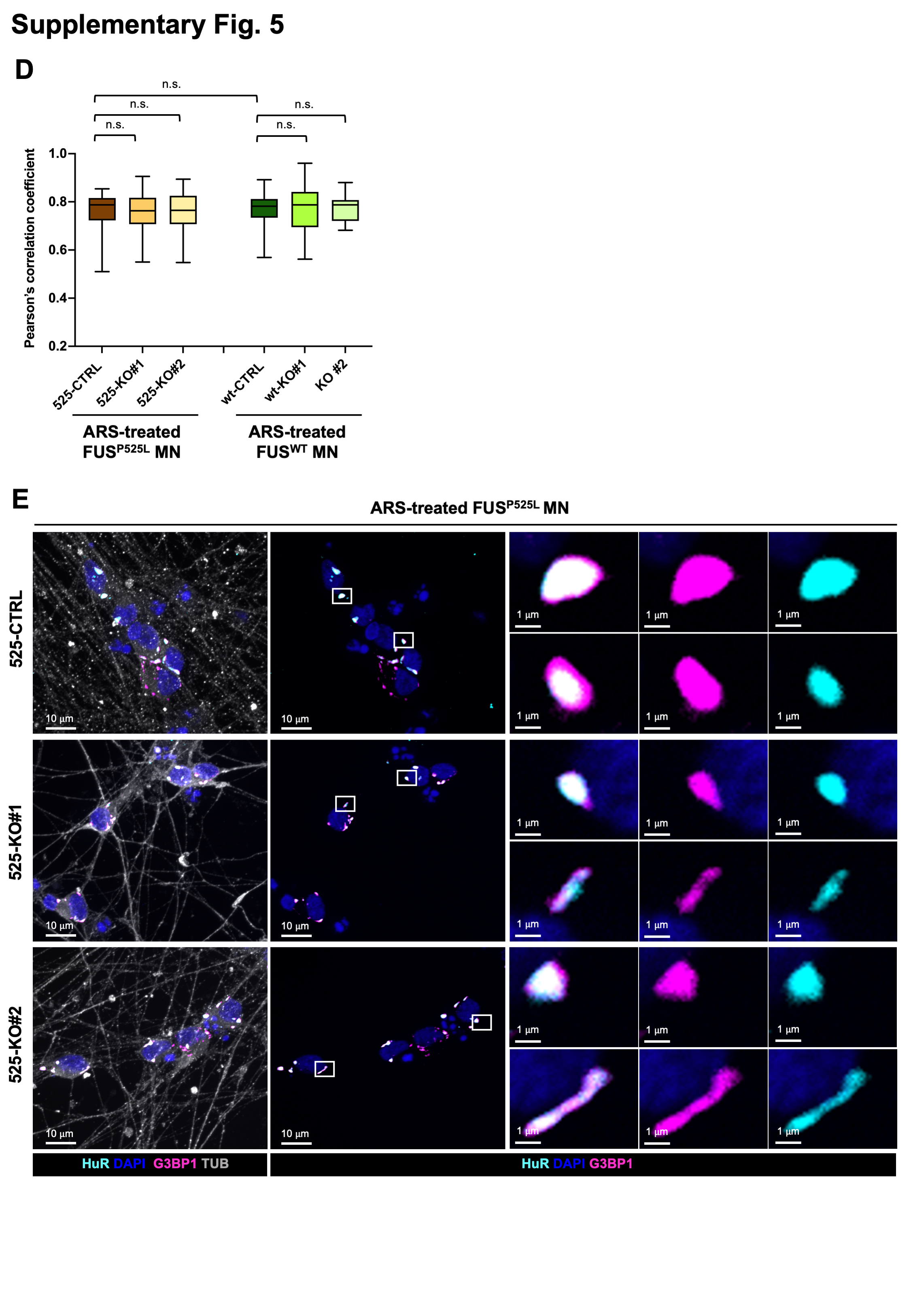

### Supplementary Fig. 5 F

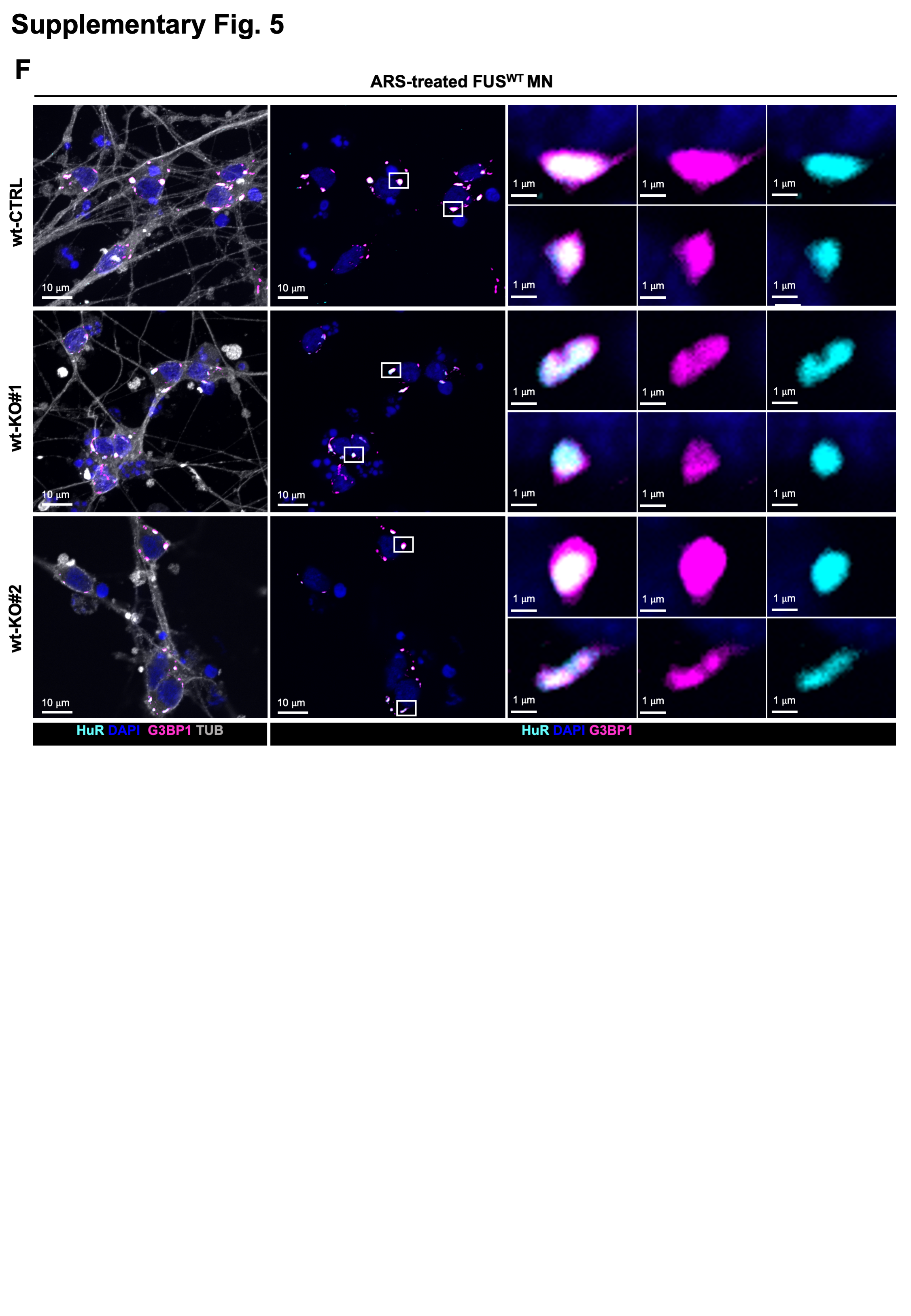
